# The ER membrane protein complex acts as a chaperone to promote the biogenesis of multi-bundle membrane proteins

**DOI:** 10.64898/2026.01.14.699575

**Authors:** Marinda Stanton, Bharti Singal, Mahamaya Biswal, Megha Agarwal, Caroline Elizabeth Scheuing, Gerardo Dasaev Vargas, Alex Gao, Casey A Gifford, Tino Pleiner

## Abstract

Nearly half of the ∼5,000 human membrane proteins need to assemble into stoichiometric complexes as part of their biogenesis at the endoplasmic reticulum (ER) membrane. How ER resident biogenesis factors coordinate membrane insertion, folding and assembly is not well understood. Here, we demonstrate that the ER membrane protein complex (EMC) insertase additionally acts as a chaperone to facilitate the assembly of heterotrimeric voltage-gated calcium channels (Ca_v_). Using function-separating mutations and inhibitory nanobodies we show that nascent Ca_v_ channels are degraded prematurely when EMC’s chaperone function is selectively perturbed. Blocking EMC’s chaperone function strongly impaired Ca_v_-dependent cardiomyocyte contraction. EMC engagement of the pore-forming Ca_v_α-subunit occurred co-translationally and required Ca_v_α’s first transmembrane domain bundle to protrude from the nascent ribosome•Sec61•multipass translocon complex. Our findings establish a chaperone function for the EMC and reveal that biogenesis of multi-bundle membrane proteins requires a highly orchestrated, co-translational interplay between ER biogenesis factors.

## INTRODUCTION

Around 25% of the human genome encodes membrane proteins that support myriad essential physiological functions – ranging from cellular nutrient uptake to signal transduction. Due to these central roles, membrane proteins are heavily overrepresented among FDA-approved drug targets (∼60%). Despite their importance for cellular physiology and human health, and intensive study over the last 50 years, fundamental aspects of membrane protein biogenesis and quality control remain poorly understood. The vast majority of membrane proteins are initially inserted into the lipid bilayer, folded and assembled into complexes at the endoplasmic reticulum (ER) membrane (Hegde and Keenan, 2024). Misfolded or unassembled membrane proteins are selectively recognized by quality control factors, ubiquitinated and extracted from the ER membrane for proteasomal degradation in the cytosol (Wu and Rapoport, 2018; Christianson and Carvalho, 2022).

An initial step in membrane protein biogenesis is the co- or post-translational insertion of the transmembrane-spanning segments (TMs) of a membrane protein into the ER lipid bilayer, which is facilitated by a limited set of ER biogenesis factors called insertases. Four insertases are known to operate at the human ER membrane and have been suggested to serve different types of clients, including the Sec61 complex (Deshaies and Schekman, 1987; Görlich et al., 1992), also called translocon, the GET1/2 complex (Schuldiner et al., 2008; Mariappan et al., 2011), the ER membrane protein complex (EMC) (Jonikas et al., 2009; Christianson et al., 2011) and the GEL complex (McGilvray et al., 2020; Sundaram et al., 2022). Recent work in the field has elucidated the structural features of these insertases (McDowell et al., 2020; McGilvray et al., 2020; Miller-Vedam et al., 2020; O’Donnell et al., 2020; Pleiner et al., 2020; Bai et al., 2020; Smalinskaitė et al., 2022) and enabled mechanistic insights into the membrane insertion of simple tail-anchored and multipass membrane proteins.

The EMC is a 324 kDa membrane protein insertase composed of nine subunits, including six core (EMC1,2,3,5,6,8) and three peripheral subunits (EMC4,7,10) (Volkmar et al., 2019). All human cells additionally express EMC9, a mutually exclusive EMC8 paralog, which was previously shown to compensate EMC8 loss and to be functionally redundant with EMC8 for EMC’s insertase function (O’Donnell et al., 2020). The EMC was shown to insert N- and C-terminal TMs of both single-pass and multipass membrane proteins that are flanked by relatively short polypeptide segments facing the ER lumen, a topology called N_exo_ / C_exo_, (Guna et al., 2018; Chitwood et al., 2018; Wu et al., 2024). Structural studies revealed that insertion occurs through a locally thinned lipid bilayer at EMC’s hydrophilic vestibule, henceforth called insertase side (Pleiner et al., 2020; Pleiner et al., 2023). The insertase side contains polar and positively charged intramembrane residues located in EMC3 and EMC6, whose mutation was shown to cause selective degradation of EMC’s insertion clients (Pleiner et al., 2020; Miller-Vedam et al., 2020; Pleiner et al., 2023).

Recently, it was proposed that the subsequent insertion of internal TMs of multipass membrane proteins occurs predominantly at a novel Sec61-containing assembly termed the multipass translocon (MPT) (McGilvray et al., 2020). When two or more TMs of a membrane protein have been translated by a ribosome docked at Sec61, the MPT assembles from the GEL, BOS and PAT complexes (Smalinskaitė et al., 2022; Sundaram et al., 2022). Cryo-electron microscopy (cryo-EM) structures of the MPT showed that these three subcomplexes enclose a small lipid-filled cavity at the back of a closed Sec61 channel. Crosslinking data using single 7xTM bundle G-protein coupled receptors as model clients suggested that the GEL and PAT complexes insert and fold internal TMs of multipass membrane proteins, respectively (Chitwood and Hegde, 2020; Smalinskaitė et al., 2022; Sundaram et al., 2022). Given the finite dimensions of the lipid-filled cavity, it remains unknown how more complex multipass membrane proteins with multiple TM bundles such as ion channels and transporters are accommodated. How folding intermediates of such proteins are protected from misfolding or the premature recognition by ER protein quality has not been explored experimentally but was suggested to require chaperones interacting with the MPT (Sundaram et al., 2025). Indeed, the EMC was recently found to interact directly with isolated BOS complex (Page et al., 2024).

In addition to membrane insertion and folding, around half of all membrane proteins additionally need to assemble into defined oligomeric assemblies to function (Juszkiewicz and Hegde, 2018). For soluble protein complexes it is well established that assembly is highly regulated and occurs either co-translationally via interactions between nascent chains or post-translationally with the help of dynamically interacting chaperones, which shield hydrophobic interaction interfaces to prevent promiscuous interactions that could lead to potentially toxic, aberrant complex or aggregate formation (Tomko and Hochstrasser, 2013; Peña et al., 2017; Shiber et al., 2018). In contrast, how hydrophilic subunit interfaces of membrane protein complexes are protected within the hydrophobic core of the lipid bilayer is not well understood. Recent data suggest that membrane protein complex assembly is indeed highly regulated and involves a growing number of membrane protein chaperones that engage and protect unassembled subunits (Gu et al., 2016; Li et al., 2017; Pleiner et al., 2021; Hooda et al., 2024).

The EMC has long been speculated to have additional functions in membrane protein biogenesis beyond its insertase role and was suggested to either act as a membrane protein chaperone itself or to serve as a recruitment platform for such chaperones (Jonikas et al., 2009; Shurtleff et al., 2018; Miller-Vedam et al., 2020; Klose et al., 2025). However, the so-far only validated direct interaction of the EMC with a potential client emerged from the cryo-EM structure of the EMC bound to a putative assembly intermediate of the cardiac voltage-gated calcium channel (Ca_v_) Ca_v_1.2 (Chen et al., 2023). Ca_v_ channels are heterotrimeric complexes composed of a pore-forming α-subunit with 24 TMs arranged in four 6-TM bundles, a cytosolic β-subunit and a lumenal α_2_δ-subunit. Purification of a heterologously expressed Ca_v_1.2α•Ca_v_β_3_ assembly intermediate strongly enriched stably bound EMC. The cryo-EM structure of this intermediate revealed that the Ca_v_1.2α•Ca_v_β_3_ complex engages EMC surfaces that are located opposite to its insertase side (**Fig. 1A**), suggesting that they could support a potential chaperone function of the EMC. Indeed, complete EMC loss was shown to reduce cell surface levels of functional Ca_v_1.2 channels (Chen et al., 2023). Building on these exciting findings, we wanted to explore whether this is due to the loss of EMC’s insertase or chaperone functions. Thus, we set out to ask if the EMC is indeed required as a chaperone for Ca_v_ assembly in human cells and to define the functional relevance of the proposed chaperones sites, as well as the timing of EMC’s engagement.

**Figure 1.**
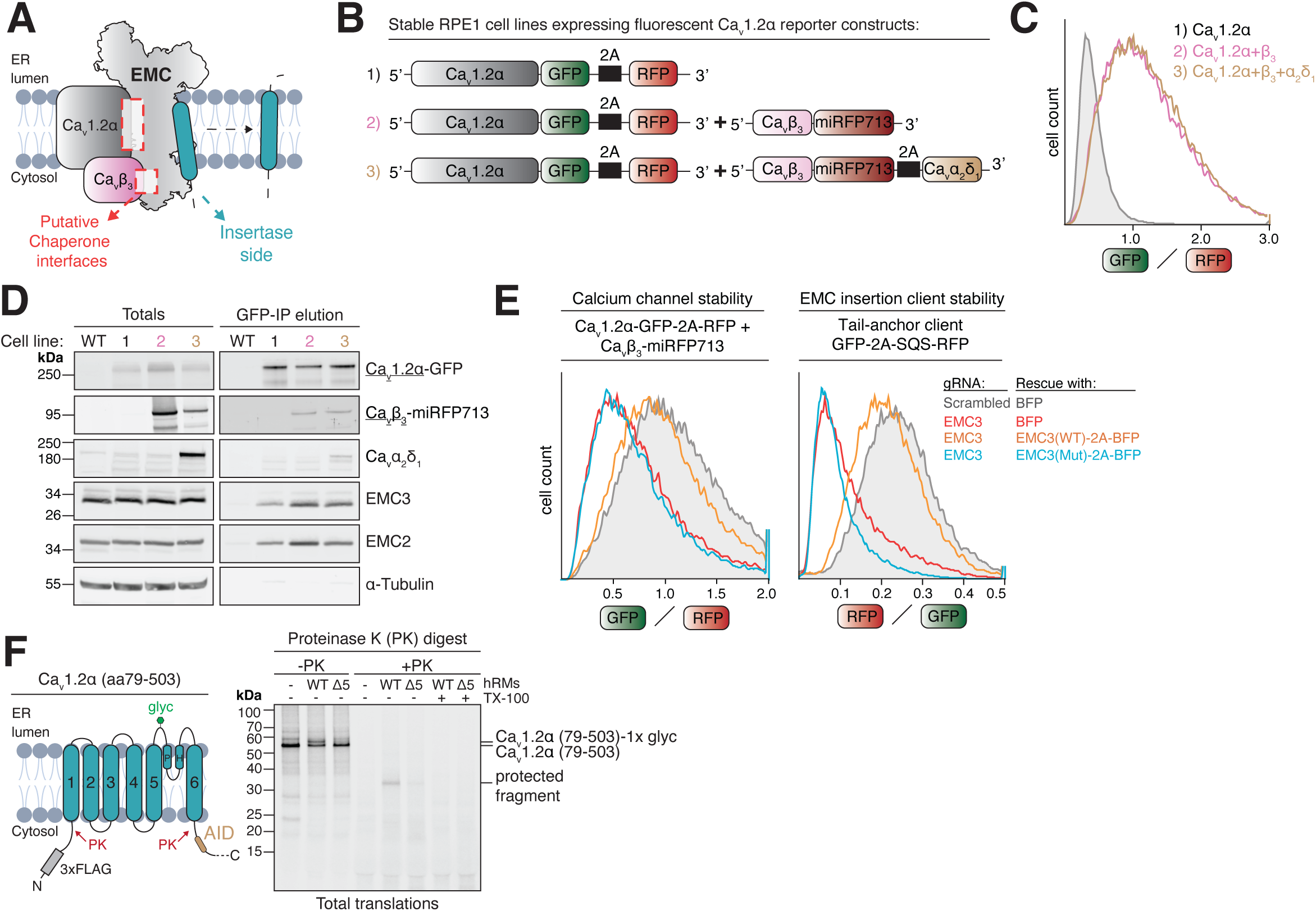
ER membrane insertion of the cardiac calcium channel Ca_v_1.2 is impaired in EMC-deficient cells. **A)** Schematic overview of the two proposed functions of the EMC in membrane protein biogenesis. The putative chaperone interfaces with the Ca_v_1.2α•β heterodimer are located on the opposite side of EMC’s well-characterized insertase side. **B)** Schematic of the lenti-viral constructs used to generate three separate stable reporter cell lines in RPE1 dCas9-BFP-KRAB cells (Jost *et al*., 2017) expressing either just Ca_v_1.2α (1), Ca_v_1.2α and Ca_v_β_3_ (2) or the full heterotrimeric Ca_v_1.2α•Ca_v_β_3_•Ca_v_α_2_δ_1_ complex (3). The GFP/RFP ratio of all cell lines reports on the post-translational stability of Ca_v_1.2α. **C)** Flow cytometry analysis of cell lines 1-3 described in B. **D)** Cell pellets of cell lines 1-3 were solubilized in detergent and subjected to anti-GFP nanobody purification of Ca_v_1.2α-GFP. Total cell lysates and protease elution of the GFP-IP were analyzed by western blotting with the indicated antibodies. **E)** Knockdown of EMC3 by CRISPRi in Ca_v_1.2α•Ca_v_β_3_ reporter cell line 2 (*left*) and RPE1 dCas9-BFP-KRAB wild-type cells transiently transduced with EMC insertase client reporter SQS (*right*). Knockdown was rescued with either just BFP, wild-type (WT) EMC3 or the insertase-deficient EMC3 R31A+R180A mutant (Mut). The GFP/RFP (Ca_v_1.2α) or RFP/GFP (SQS) ratios of BFP positive cells were determined by flow cytometry and are depicted as histograms. **F)** Insertion defect of Ca_v_ 1.2α in EMC5 KO (Δ5) ER membranes. ^35^S-methionine labeled Ca_v_ 1.2α (amino acids [aa] 79-503), a fragment corresponding to a 6xTM domain comprising the first voltage-sensing and pore domain, carrying an N-terminal 3xFLAG tag was *in vitro* translated in rabbit reticulocyte lysate supplemented with human rough ER membranes (hRMs). Non-incorporated as well as cytosolically accessible protein portions were digested with proteinase K (PK) in the presence or absence of Triton-X-100 (TX-100) to solubilize hRMs. The resulting protease protected fragment (PF) is highlighted and is substantially reduced in Δ5 hRMs compared to wild-type (WT) hRMs. PH = pore helix. AID = alpha-interacting domain. glyc = glycosylation.

## RESULTS

### Voltage-gated calcium channel biogenesis is sensitive to loss of EMC’s insertase activity

To investigate EMC’s role in the biogenesis of voltage-gated calcium channels we created a series of stable fluorescent reporter cell lines to assess how mutations at the opposing chaperone and insertase sides of the EMC (**Fig. 1A**) affect calcium channel stability. All cell lines express Ca_v_1.2α fused to a C-terminal EGFP and a separate, cytosolic RFP as a translation normalization control (**Fig. 1B**). The GFP/RFP ratio distribution, determined by flow cytometry, reports on the post-translational stability of the α-subunit under different conditions (**Fig. 1C**). Additional stable cell lines further express either the auxiliary cytosolic Ca_v_β_3_-subunit fused to far-red fluorescent protein miRFP713 (Matlashov et al., 2020) or the entire heterotrimeric channel, including the lumenal Ca_v_α_2_δ_1_ subunit. We validated that all proteins were expressed and formed a complex (**Fig. 1D**). As previously reported, expression of the β-subunit strongly stabilized the α-subunit (Altier et al., 2011; Waithe et al., 2011), as indicated by an increase in the GFP/RFP ratio. Additional expression of Ca_v_α_2_δ_1_ had no further stabilizing effects, and as previously shown, reduced EMC co-purification with the α-subunit (**Fig. 1C-D**), likely due to a steric clash between their binding sites (Chen et al., 2023). Knockdown or knockout of core EMC subunits reduced total EMC levels in cells and led to strong degradation of the α-subunit, as evident from a decrease in the GFP/RFP ratio (**Fig. S1A**) and a visible reduction of full-length channel levels by western blot (**Fig. S1B**). In both cases, normal levels of the channel could be restored by treating cells with the proteasome inhibitor bortezomib. As expected, bortezomib-stabilized channels were highly ubiquitylated (**Fig. S1B**). This indicates that the α-subunit is degraded by the ubiquitin-proteasome system in the absence of the EMC, suggesting a critical role for the EMC in its membrane insertion, folding or assembly.

To test the reliance of Ca_v_1.2α on EMC’s insertase function, we introduced a well-characterized double mutation of two key Arginine residues (R31A+R180A) into EMC3 (Pleiner et al., 2023). Wild-type EMC3 add-back to EMC3 CRISPRi knockdown cells rescued the stability of EMC insertase clients SOAT1 and SQS. As expected, add-back of the R31A+R180A double mutant strongly destabilized both clients, but had no effect on the GET1/2 client VAMP2 (**Fig. 1E, Fig. S1C-D**). Given the absence of terminal TMs with N_exo_ or C_exo_ topology, it was surprising that the calcium channel behaved like an EMC insertion client and was degraded when EMC3’s R31 and R180 residues were mutated (**Fig. 1E**). To test if EMC loss affects insertion of newly synthesized α-subunits into the ER membrane more directly, we used protease protection assays to probe the amount of *in vitro* translated channel that was inserted in the correct topology in wild-type versus EMC5 KO ER membranes (**Fig. 1F, Fig. S1E-G**). Mirroring our flow cytometry data, we found a striking defect in the insertion of the first 6-TM bundle of the α-subunit in EMC5 KO ER membranes, whereas a non-EMC dependent control membrane protein was unaffected (**Fig. 1F, Fig. S1F-G**). Given the lack of canonical insertase client features, the α-subunit might represent a non-canonical insertase client or its insertion may be indirectly affected by EMC loss. Any attempt to validate a potential chaperone function of the EMC thus needs to include controls showing that its insertase function is left intact. This precludes the use of conditions that lead to complete EMC loss, such as the knockout of a core EMC subunit. We thus set out to generate mutations and inhibitors that could selectively perturb EMC’s potential chaperone activity without affecting its insertase activity.

### Function-separating mutations establish that the EMC acts as a chaperone in calcium channel biogenesis

The EMC•Ca_v_1.2α•Ca_v_β_3_ cryo-EM structure identified two interaction interfaces, an intramembrane interface between Ca_v_1.2α and EMC1, and a cytosolic interface between Ca_v_β_3_ and EMC8 (**Fig. 2A**) (PDB: 8EOI; Chen et al., 2023). To directly test if these interfaces are important for calcium channel biogenesis in human cells, we sought to introduce mutations on EMC1 and EMC8 at these surfaces. Starting with the cytosolic interface, we depleted EMC8 and its mutually exclusive paralog EMC9 using specific siRNAs and assessed calcium channel stability (**Fig. S2A-E**). Complete EMC loss resulting from double knockdown of both EMC8 and EMC9 strongly destabilized Ca_v_1.2α (**Fig. S2A-B**). Strikingly, single knockdown of EMC8, but not EMC9, was sufficient for Ca_v_1.2α destabilization (**Fig. S2B**). This was surprising since EMC8 and EMC9 were previously found to be functionally redundant for insertase client reporters of the EMC (O’Donnell et al., 2020). Our data confirmed that insertase clients were not affected by single knockdowns of EMC8 or EMC9 but did respond to double knockdown of both subunits (**Fig. S2E**). Closer inspection by western blotting, however, showed that EMC8 knockdown in both RPE1 and HEK293 cells, as well as EMC8 knockout in HEK293 cells, resulted in substantially reduced total EMC levels and thus was not fully compensated by EMC9 upregulation (**Fig. S2B-D**).

**Figure 2.**
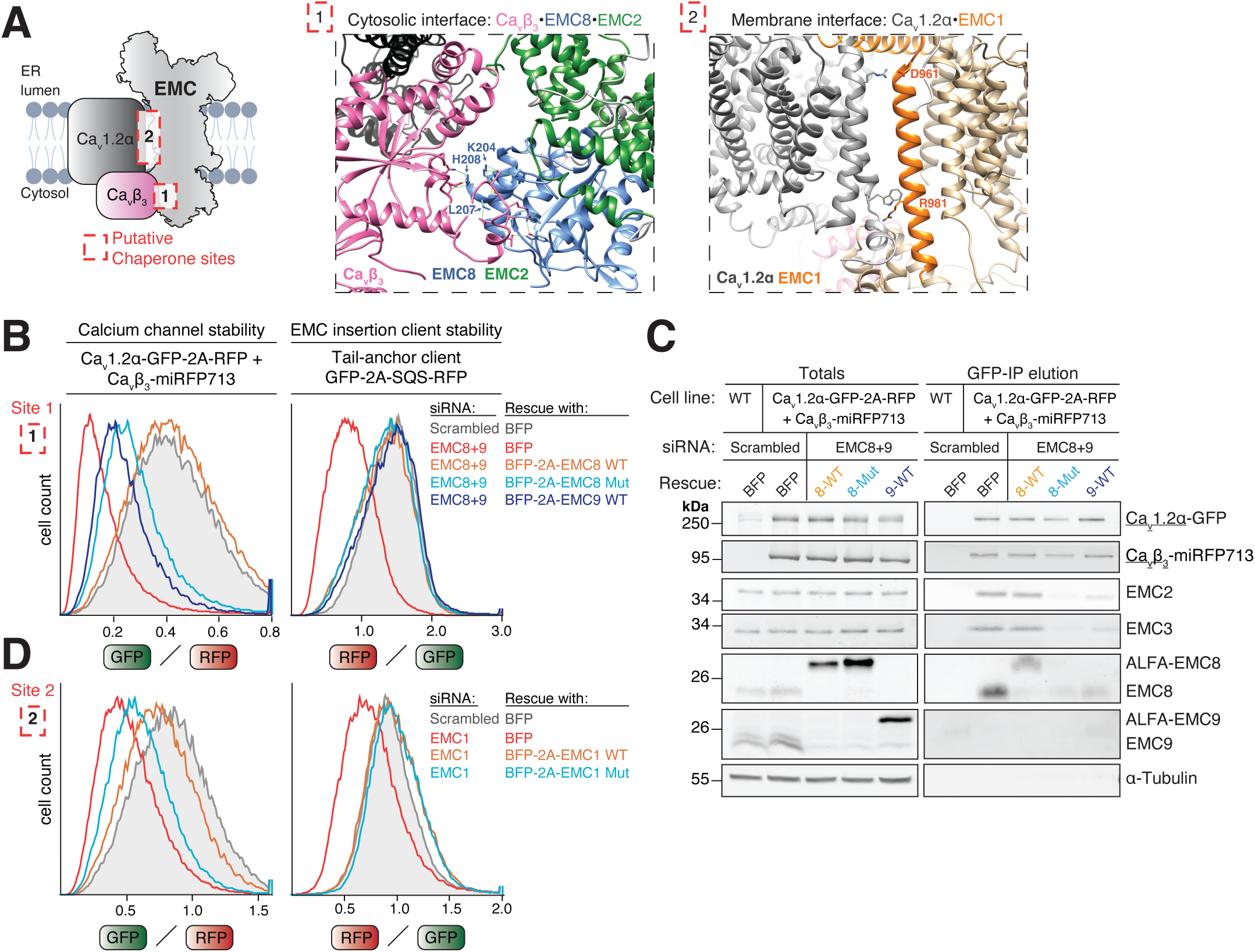
EMC’s intramembrane and cytosolic chaperone interfaces are required for Ca_v_1.2 calcium channel biogenesis in cells. **A)** Detailed views of the cytosolic interface between Ca_v_β_3_ and EMC8 [site 1] and the intramembrane interface between Ca_v_1.2α and EMC1 [site 2] based on PDB 8EOI (Chen *et al*., 2023). Critical interface residues on EMC8 and EMC1 mutated below are highlighted as sticks. **B)** 72h scrambled or EMC8+9 double siRNA knockdown in stable RPE1 Ca_v_1.2α•Ca_v_β_3_ reporter cells or RPE1 wild-type (WT) cells transiently transduced with EMC-dependent insertase client reporter SQS. 24h after siRNA transfection, cells were transduced with lenti-viral rescue constructs encoding either just BFP or BFP separated by a 2A site from either EMC8 WT, EMC8 K204A, L207A, H208A mutant (Mut) or EMC9 WT. The GFP/RFP (Ca_v_1.2α) or RFP/GFP (SQS) ratios of BFP positive cells were determined by flow cytometry and are depicted as histograms. **C)** siRNA knockdown and rescue assays in stable RPE1 Ca_v_1.2α•Ca_v_β_3_ reporter cells were performed as described in B. Cells were lysed in detergent and subjected to anti-GFP nanobody purification of Ca_v_1.2α-GFP to assess EMC co-purification. Total cell lysates and protease elution of the GFP-IP were analyzed by western blotting with the indicated antibodies. **D)** 72h scrambled or EMC1 siRNA knockdown in stable RPE1 Ca_v_1.2α•Ca_v_β_3_ reporter cells or RPE1 wild-type (WT) cells transiently transduced with EMC-dependent insertase client reporter SQS. 24h after siRNA transfection, cells were transduced with lenti-viral rescue constructs encoding either just BFP or BFP separated by a 2A site from either EMC1 WT or EMC1 D961A, R981L mutant (Mut). Analysis by flow cytometry as in B.

To rule out the possibility that lower total levels of EMC, and the associated reduction of insertase activity, caused calcium channel sensitivity to EMC8 loss, we turned to an siRNA rescue assay. After careful inspection of the cryo-EM structure (PDB: 8EOI; Chen et al., 2023), we picked three key residues on EMC8 (K204, L207 and H208) that contact Ca_v_β_3_ (**Fig. 2A**). To assess the impact of mutating these residues on calcium channel stability, we knocked down both EMC8 and EMC9 and added back siRNA-resistant wild-type or mutant EMC8. As expected, wild-type EMC8 completely rescued total EMC levels and stabilized the calcium channel and all known classes of insertase clients (**Fig. 2B, Fig. S2F-G**), highlighting the specificity of the used siRNAs. Add-back of a triple Alanine mutant EMC8 (K204A, L207A, H208A) also rescued total EMC levels and stabilized all insertion clients, indicating that these mutations do not interfere with EMC assembly or insertase activity. However, this mutant strongly destabilized Ca_v_1.2α (**Fig. 2B**). A similar magnitude of destabilization was also observed in cell lines additionally expressing the lumenal α_2_δ_1_ subunit (**Fig. S2G**), indicating that EMC dependence cannot simply be overcome by overexpressing the missing binding partner. To assess if cells expressing solely EMC9 can support Ca_v_ channel biogenesis, we rescued EMC8+9 double knockdown by add-back of siRNA-resistant EMC9. Confirming our single knockdown results, EMC9 add-back essentially phenocopied EMC8 mutant add-back and led to calcium channel degradation (**Fig. 2B**). All knockdowns and add-back conditions did not affect EMC-independent control membrane proteins VAMP2 and ASGR1 (**Fig. S2G**). To explore if mutation of the β-subunit interface on EMC8 or its replacement with EMC9 could affect interaction of the EMC with Ca_v_1.2α•β, we purified the channel and assessed EMC co-purification (**Fig. 2C**). We found that EMC complexes containing mutant EMC8 or EMC9 failed to co-purify with the calcium channel assembly intermediate, indicating that the loss of their interaction causes channel degradation in cells. This provides first evidence that the chaperone function of the EMC is indeed required for calcium channel stability in cells and further indicates that the EMC acts as a chaperone prior to the final heterotrimer assembly step. Our data reveal that the paralogs EMC8 and EMC9 are not functionally redundant for EMC’s chaperone function and suggest that specialized EMC complexes co-exist in mammalian cells.

We next wanted to evaluate if the intramembrane interface between EMC1 and Ca_v_1.2α is similarly required for Ca_v_ channel stability (**Fig. 2D**). We identified two charged residues on EMC1, D961 and R981, that we mutated to Ala and Leu, respectively. Using an siRNA rescue assay, we could show that the resulting double mutant incorporated efficiently into EMC complexes (**Fig. S2H**) and selectively destabilized the calcium channel, but not an EMC insertase client or control membrane proteins (**Fig. 2D, Fig. S2I**). These experiments validate that the EMC uses both its cytosolic and intramembrane interfaces to prevent degradation of a heterodimeric Ca_v_1.2α•β assembly intermediate and establish function-separating mutations that selectively impair EMC’s novel chaperone role. When EMC’s chaperone interfaces are mutated, Ca_v_1.2α is degraded even in the presence of an excess of otherwise stabilizing β-subunit (**Fig. 2B**, **Fig. 2D**) (Altier et al., 2011; Waithe et al., 2011), suggesting that the EMC requirement lies upstream of Ca_v_1.2α•β dimer formation.

### Selective inhibitory nanobodies validate Ca_v_ channel dependence on EMC’s chaperone function

The siRNA rescue assays described above require efficient knockdown of endogenous EMC subunits and their replacement with ectopically overexpressed copies, limiting their usefulness to cell lines that are amenable to such manipulations and potentially introducing unwanted side-effects. To provide easily portable, genetically encoded inhibitors, we sought to generate single-domain antibodies, called nanobodies (Muyldermans, 2013), targeting the endogenous EMC. We immunized an alpaca with purified EMC complex reconstituted into proteoliposomes and selected nanobodies by phage display (Pleiner et al., 2015) **(Fig. 3A)**. We then grouped nanobodies into unique classes based on their antigen-binding loop sequences (**Fig. S3A**), expressed them in *E. coli* and characterized their specificity by affinity purification from human cell lysate (Stevens et al., 2024). We provide the coding sequences of 16 distinct nanobody classes (**Table S1**) that we found specifically enriched the assembled EMC complex with high affinity and in high purity (**Fig. 3B, Fig. S3A-B**).

**Figure 3.**
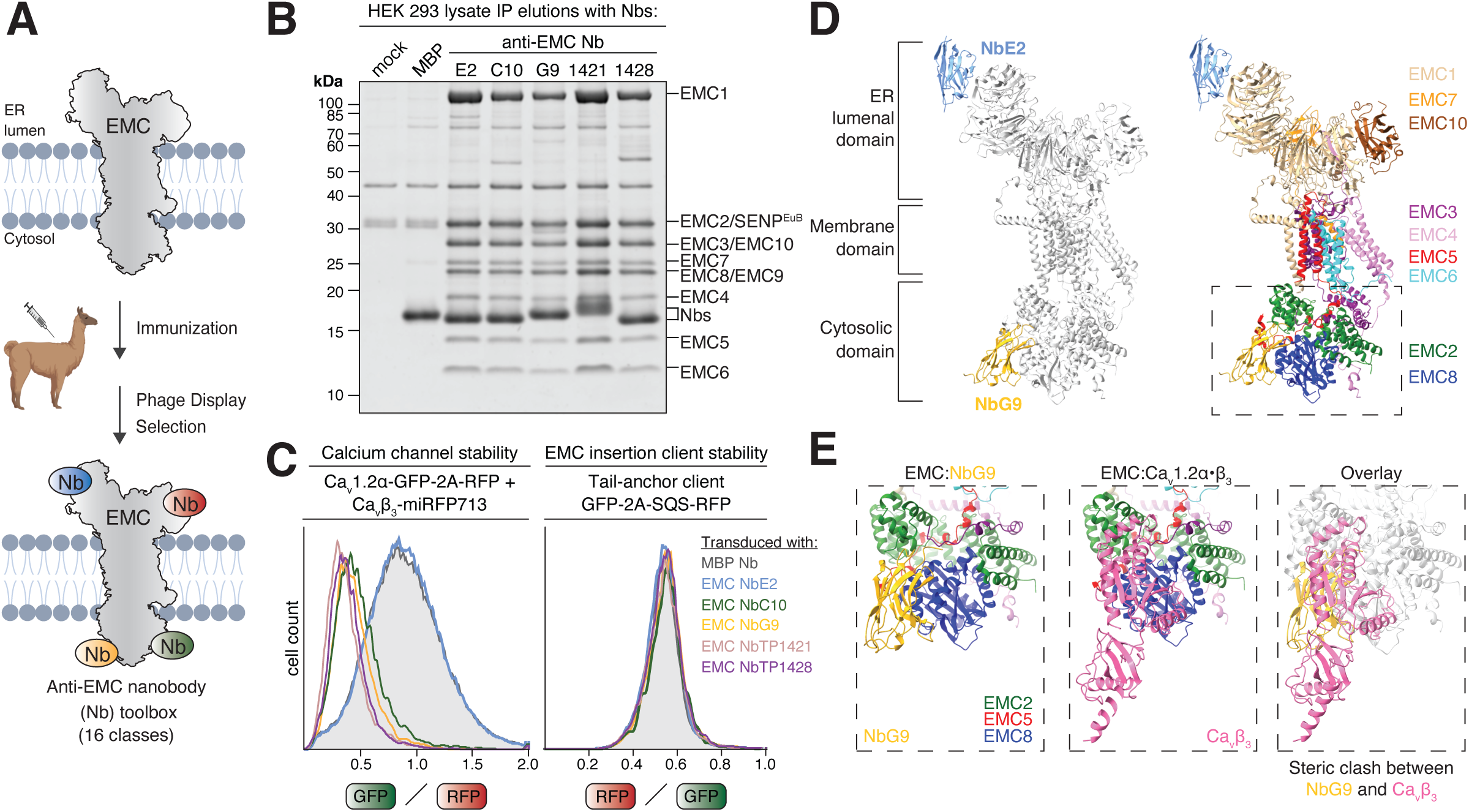
An anti-EMC nanobody toolbox yields genetically encoded inhibitors of EMC’s chaperone function. **A)** Schematic illustrating the basic workflow of nanobody (Nb) generation. Purified EMC stabilized in LMNG detergent micelles or proteoliposomes was used to immunize an alpaca. Nb coding sequences were cloned from B-lymphocytes of a small blood sample and the resulting Nb library screened by phage display to identify anti-EMC Nbs. 16 different classes were characterized. **B)** Biotinylated, SENP^EuB^ protease-cleavable anti-EMC Nbs or an anti-Maltose-binding protein (MBP) control Nb were immobilized on Streptavidin magnetic beads and incubated with a GDN-solubilized Expi293 total cell lysate. After washing, Nbs along with their bound proteins were eluted by native SENP^EuB^ cleavage. The eluate was analyzed by SDS-PAGE and SYPRO Ruby staining. Note that all anti-EMC Nbs specifically purify all nine known EMC subunits. **C)** Flow cytometry assay as in Fig. 2B, but with lentiviral transduction of the indicated Nbs, expressed from a BFP-2A-Nb cassette. The GFP/RFP or RFP/GFP ratios of BFP-positive cells are depicted as histograms. Note that anti-EMC Nbs G9, C10, TP1421 and TP1428 selectively inhibit EMC’s chaperone, but not insertase activity. **D)** Model of the human EMC in complex with inert NbE2 and inhibitory NbG9 determined using cryo-electron microscopy (cryo-EM) (PDB 9ZZ6). **E)** Views of EMC’s cytosolic chaperone interface formed by EMC2 and EMC8 in complex with NbG9 (*left*, our structure), in complex with Ca_v_β_3_ (*middle*, PDB 8EOI, Chen et al., 2023) or an overlay of both (*right*) to highlight that binding of NbG9 sterically blocks Ca_v_β_3_ binding to the EMC.

We were curious to explore if intracellular expression and binding of these nanobodies to the endogenous EMC could interfere with its chaperone or insertase function. Given that these nanobodies might potentially bind to either EMC’s lumenal or cytosolic domains, we expressed each nanobody in both compartments, alongside fluorescent reporters of chaperone or insertase function (**Fig. S3C**). Systematic screening revealed four distinct classes of nanobodies that selectively inhibited EMC’s chaperone function without affecting its insertase function in cells. This was demonstrated by the strong degradation of Ca_v_1.2α, while the biogenesis of the insertase client SQS remained essentially unimpaired (**Fig. 3C**). The expression of these four inhibitory nanobodies in cells also substantially reduced EMC co-purification with Ca_v_1.2α•β (**Fig. S3D**) and essentially phenocopied our interface mutants. These findings suggest that binding of these nanobodies either sterically or allosterically interfered with binding of Ca_v_1.2α•β to the EMC.

To obtain structural insights into the inhibition mechanism, we purified endogenous EMC using the inert control nanobody (Nb) NbE2 and formed a complex with inhibitory NbG9 (**Fig. S4A-B**). The resulting EMC•NbE2•NbG9 complex migrated as a single peak on size-exclusion chromatography and was subjected to structural analysis by cryo-EM (**Fig. 3D-E, Fig. S4C-F, Table S2**). Our nanobody co-structure (PDB 9ZZ6) revealed the inert NbE2 bound to the lumenal top β-propeller domain of EMC1, whereas inhibitory NbG9 was bound to EMC’s cytosolic domain. Closer inspection of the NbG9 binding site showed that it contacts both EMC2 and EMC8, and comparison with the EMC•Ca_v_1.2α•β structure suggested that binding of NbG9 sterically clashes with binding of the Ca_v_β-subunit to the EMC. Interface mutations in both NbG9 and EMC8 validated this structural model (**Fig. S5**). These findings indicate that NbG9 inhibits EMC’s chaperone function by directly blocking its cytosolic chaperone interface.

In summary, we have created a toolbox of EMC-specific nanobodies that can be used to purify the endogenous human EMC under native conditions (Stevens et al., 2024). Four of these nanobody classes represent selective, genetically encoded inhibitors of EMC’s chaperone function that act on the endogenously expressed EMC and so provide orthogonal validation of its importance for calcium channel biogenesis without the need to knockdown and replace EMC subunits. These nanobodies can now easily be introduced into various human cell lines to study the physiological relevance and client spectrum of this novel, still vastly underexplored role of the EMC in membrane protein biogenesis.

### Neuronal and skeletal muscle calcium channels also depend on EMC chaperone function

We further sought to establish if EMC’s chaperone function extends to related voltage-gated calcium channels expressed in the human brain and skeletal muscle. We created stable fluorescent reporter cell lines for the brain-specific P/Q-type channel Ca_v_2.1 and the skeletal muscle-specific L-type channel Ca_v_1.1, both co-expressed with the Ca_v_β_3_ subunit. Using both function-separating mutations and inhibitory nanobodies, we could reveal a similar dependence of these channels on EMC’s cytosolic chaperone interface for stability in cells (**Fig. 4A, Fig. S6A**). Additionally, we wanted to test if all four human Ca_v_β-subunit paralogs support EMC engagement and created stable cell lines expressing these subunits alongside Ca_v_1.2α. Indeed, we found that all four β-subunits strongly stimulated EMC binding to Ca_v_1.2α (**Fig. S6B**). Collectively, these results, in combination with overall high sequence conservation of EMC’s binding sites on Ca_v_1 and Ca_v_2 family α-subunits (Chen et al., 2023), strongly suggest that EMC’s chaperone role extends to voltage-gated calcium channels expressed throughout the human body.

**Figure 4.**
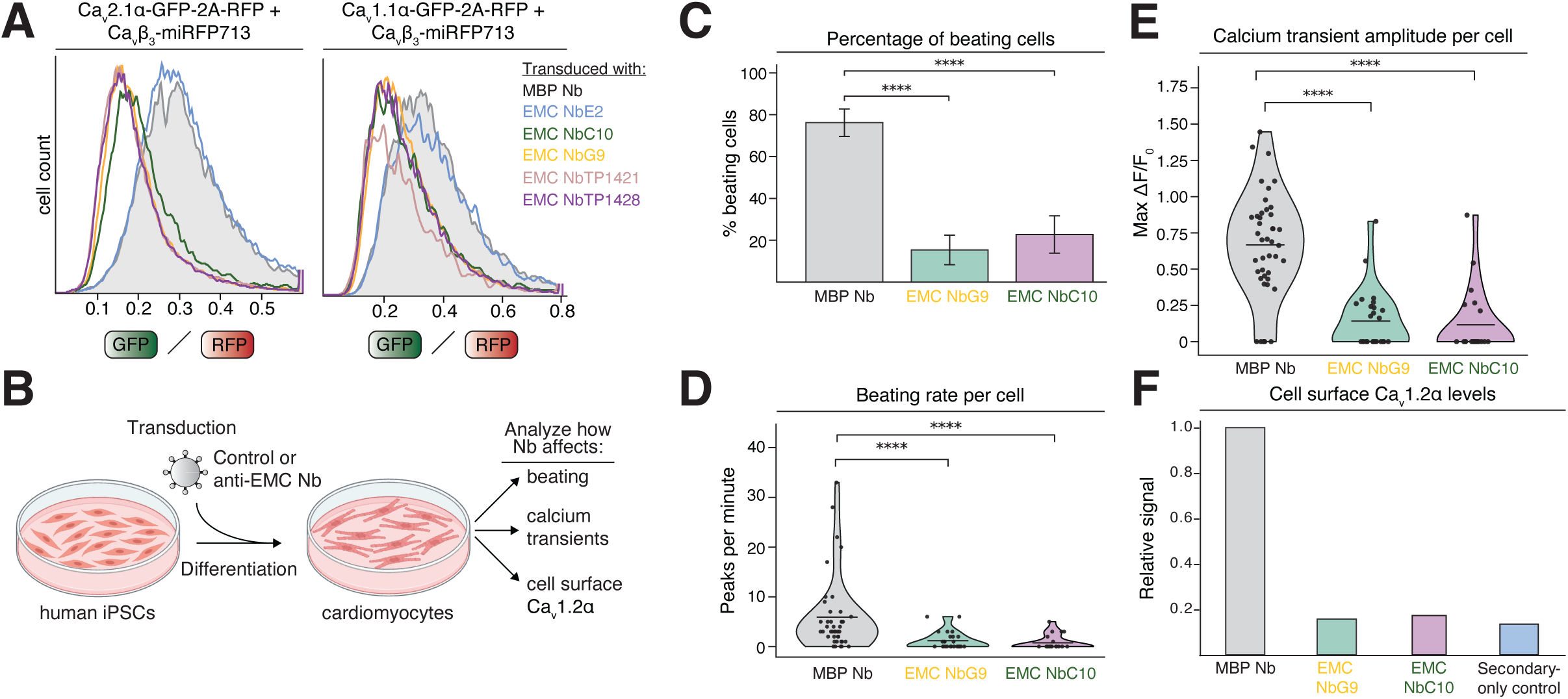
Blocking EMC’s chaperone function impairs cardiomyocyte contraction. **A)** Experiments in stable RPE1 cells expressing the brain Ca_v_2.1α•Ca_v_β_3_ channel (*left*) or skeletal muscle Ca_v_1.1α•Ca_v_β_3_ channel reporter (*right*). Cells were transduced to express the indicated control or inhibitory nanobodies (Nb) from a BFP-2A-Nb cassette in the cytosol. The GFP/RFP ratios of BFP^+^ cells are depicted as histograms. **B)** Experimental workflow schematic. Human iPSCs were transduced with control or anti-EMC nanobodies, differentiated into cardiomyocytes and then the depicted traits were analyzed. **C)** Bar plot showing the percentage of beating BFP^+^ and GCaMP^+^ cardiomyocytes transduced with either control anti-MBP Nb, anti-EMC NbG9 or C10. Values represent mean ± standard error. p < 0.0001. **D)** Violin plot showing the distribution of beating rates (peaks per minute) of BFP^+^ and GCaMP^+^ cardiomyocytes transduced with either control anti-MBP Nb, anti-EMC NbG9 or C10. Each point represents an individual cell. Mean values are indicated by horizontal bars. p < 0.0001. **E)** Violin plot showing the distribution of calcium transient amplitudes (max ΔF/F0) in BFP^+^ and GCaMP^+^ cardiomyocytes transduced with either control anti-MBP Nb, anti-EMC Nb G9 or C10. Mean values are indicated by horizontal bars. Each point represents an individual cell. p < 0.0001. **F)** Staining of GCaMP^+^ cardiomyocytes transduced with either control anti-MBP Nb, anti-EMC NbG9 or C10 under non-permeabilizing conditions using an antibody targeting an extracellular epitope on Ca_v_1.2α to detect Ca_v_1.2α’s cell surface levels. The primary antibody was detected with an Alexa 647-labeled secondary antibody. The median Alexa 647 signal of BFP^+^ cells is plotted. A secondary antibody-only staining was included as a control for background fluorescence.

### Blocking EMC’s chaperone function impairs cardiomyocyte contraction

Collectively, our data using exogenously expressed calcium channel reporters in human cell lines indicate that loss of EMC chaperone function leads to channel degradation. To explore if blocking EMC chaperone function would impact the function of an excitable cell type that endogenously expresses the Ca_v_1.2 channel, we turned to human induced pluripotent stem cell (iPSC)-derived cardiomyocytes (CMs) as a model system (**Fig. 4B**). We transduced differentiating cardiomyocytes expressing the fluorescent calcium indicator GCaMP6f (Huebsch et al., 2015) with either anti-MBP control or inhibitory anti-EMC nanobodies (Nbs) G9 or C10 and analyzed both the percentage of beating CMs, as well as the beating rate per cell from videos recorded by live-cell imaging (**Fig. 4C-D, Video S1-3**). Both parameters were dramatically and selectively reduced upon expression of both inhibitory Nbs. We further directly quantified calcium transients associated with CM contraction. CMs transduced with control Nb showed nicely synchronized, circulating calcium waves accompanying contraction, whereas inhibitory Nb-treated cells showed very little GCaMP activation (**Fig. 4E**). These findings highlight that blocking EMC’s chaperone function dramatically reduces CM contraction and identify altered calcium dynamics as an underlying cause. To directly assess the amount of Ca_v_1.2 channel expressed at the cell surface in CMs treated with control or inhibitory nanobodies, we stained non-permeabilized CMs with an anti-Ca_v_1.2α antibody that recognizes an extracellular epitope (**Fig. 4F**). Indeed, we found that blocking EMC chaperone function reduces plasma membrane Ca_v_1.2α levels, suggesting that Ca_v_1.2 channel loss contributes to the lack of calcium transients and thus strongly reduces CM contraction.

These data provide first indications that loss of EMC’s chaperone activity impairs the function of an important excitable cell type and suggest that its loss cannot easily be compensated by other chaperones.

### The EMC co-translationally engages nascent α-subunits at the multipass translocon

While the cryo-EM structure of the EMC•Ca_v_1.2α•β complex revealed the EMC bound to full-length Ca_v_1.2α, it also indicated that the EMC predominantly engages the N-terminal TM bundle I that would emerge first during Ca_v_1.2α’s co-translational membrane insertion (Chen et al., 2023). Our mutational data further indicated destabilization of Ca_v_1.2α by loss of EMC chaperone function even in the presence of both overexpressed partner subunits and suggested that Ca_v_1.2α’s EMC requirement precedes complex assembly. Collectively, these observations raised the possibility that EMC binding could be an early event in calcium channel biogenesis.

To probe if the EMC can engage nascent Ca_v_1.2α co-translationally, we employed a cell-free *in vitro* translation and ER membrane insertion system based on rabbit reticulocyte lysate that allows stalling membrane protein biogenesis at desired nascent chain lengths by using truncated mRNAs without stop codon (**Fig. 5A-B**) (Sharma et al., 2010). This results in nascent chains stably attached to ribosomes via peptidyl-tRNA bonds and allows arresting biogenesis intermediates for analysis of their co-translational interactome. Using this approach we generated stalled ribosome-nascent chain complexes (RNCs) exposing either 2, 4, 6 or 12xTM segments of Ca_v_1.2α. Both 6- and 12-TM stall constructs also include the binding site for Ca_v_β (Pragnell et al., 1994; Witcher et al., 1995). Membrane-associated RNCs were purified via an N-terminal ALFA-tag on Ca_v_1.2α and the eluates were pelleted through a sucrose cushion to enrich the ribosome-bound fraction. Western blot analysis confirmed that purified RNCs contained ribosomes, Sec61 and the expected tRNA-associated nascent chains (**Fig. 5C-D, Fig. S7A**). Purified Ca_v_β, supplemented into the translation reaction, associated with RNC•MPT complexes containing its binding site, suggesting that co-translational Ca_v_α•β complex formation is possible. We were further able to confirm the co-translational assembly of the MPT, as indicated by the presence of BOS complex subunit NOMO2, once two or more TM segments of Ca_v_1.2α were inserted into the membrane. Importantly, we were able to detect stable EMC recruitment when at least two TM segments of Ca_v_1.2α were membrane-inserted, strongly suggesting that it can co-translationally engage RNC•MPT complexes. While the amount of EMC engagement was constant with RNCs containing 2, 4 and 6xTMs (**Fig. S7A**), its increased abundance with RNCs containing an additional 6-TM bundle (12xTMs) (**Fig. 5C-D**), suggests that nascent chains might need to be of sufficient length to protrude from the MPT’s lipid-filled cavity to allow efficient EMC engagement.

**Figure 5.**
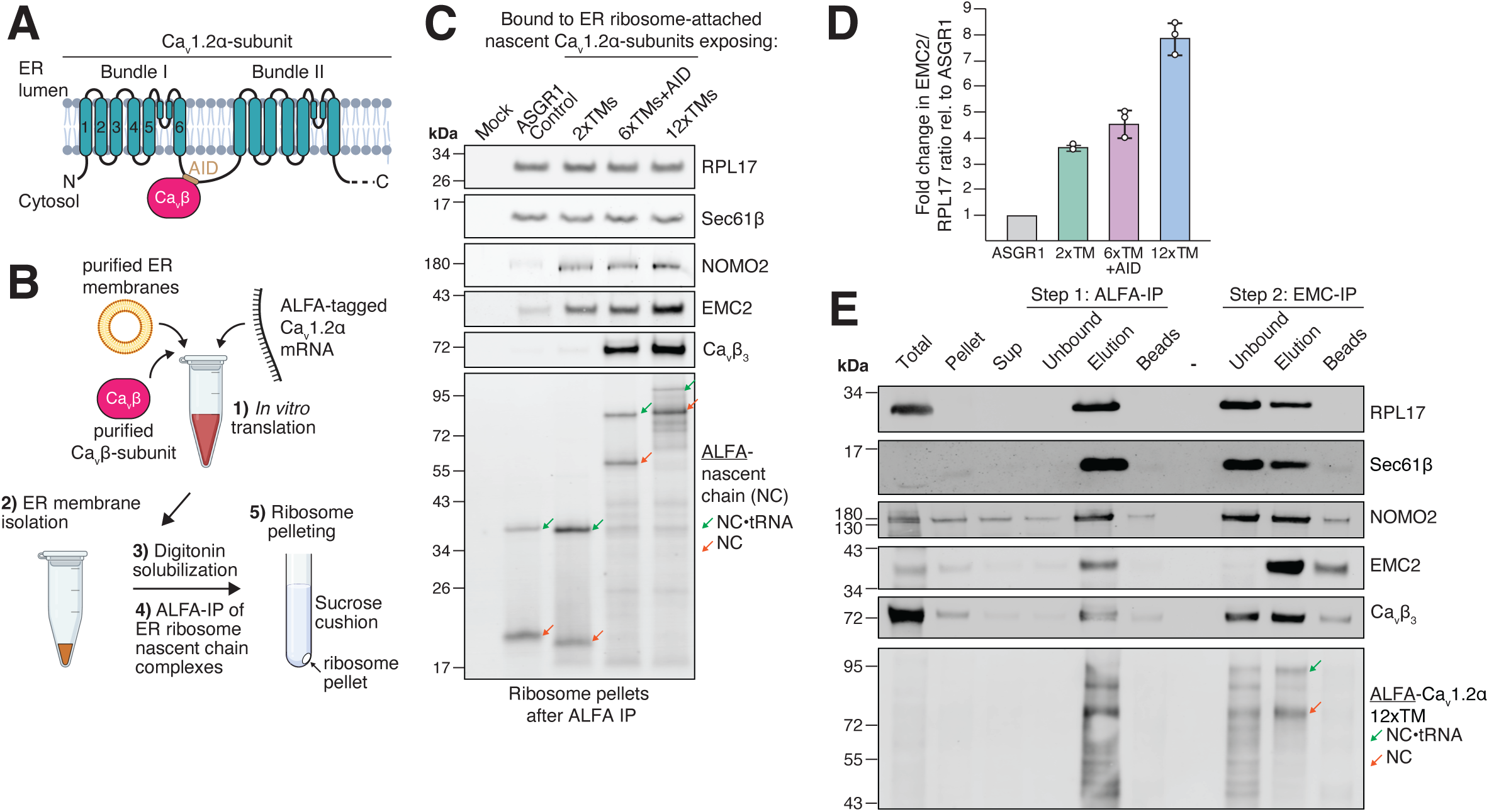
*In vitro* translation of Ca_v_1.2α reveals co-translational association with EMC and Ca_v_β_3_. **A)** Schematic illustrating the topology of Ca_v_1.2α. For clarity only two out of four 6xTM bundles are depicted. The Ca_v_β binding site, called alpha-interacting domain (AID), is located in the cytosolic linker between bundles I and II. **B)** Set-up of the cell-free *in vitro* translation and ER insertion system. Homemade, micrococcal nuclease-treated rabbit reticulocyte lysate is supplemented with purified ER membranes prepared from HEK293 cells and *in vitro* transcribed Ca_v_1.2α mRNA truncated at various strategic points to expose a desired number of TMs to the ER membrane after insertion. Ca_v_β_3_ purified from *E. coli* was supplemented as well. After *in vitro* translation, ER membranes are pelleted through a sucrose cushion, solubilized in Digitonin and ER-bound ribosome nascent chain complexes (RNCs) are purified by ALFA nanobody IP and native SUMOstar protease elution. The eluate is then spun through a second sucrose cushion to isolate the ribosome-bound fraction for analysis by western blot. **C)** Normalized ribosome pellets of RNCs stalled while translating either an EMC-independent single-pass control protein ASGR1, or Ca_v_1.2α exposing the indicated number of TMs outside the ribosome exit tunnel, were analyzed by Western blotting with the indicated antibodies. A mock translation without RNA served as a background control. Note that both EMC and Ca_v_β_3_ associate with co-translational biogenesis intermediates of Ca_v_1.2α. **D)** Quantification of the EMC2/RPL17 ratio from western blots depicted in C) for all Ca_v_1.2α stall lengths normalized to ASGR1. Error bars represent mean and standard deviation from three independent replicates. Note the increase in EMC association when more than 6xTMs of Ca_v_1.2α are membrane-inserted. **E)** Stalled RNCs containing Ca_v_1.2α exposing 12xTMs were generated as described in B. After purification of ALFA-tagged RNCs using a SUMOstar-cleavable anti-ALFA-nanobody, the resulting protease eluate was subjected to a second IP with SENP^EuB^-cleavable anti-EMC NbE2. Samples of this 2-step IP were analyzed by western blotting with the indicated antibodies. Note that the EMC is bound to intact RNCs containing tRNA-associated nascent chain (NC), ribosomes (RPL17), the translocon (Sec61β) and MPT BOS complex subunit NOMO2. EMC-IP also specifically enriches full length nascent chains.

These findings indicated that the EMC might engage nascent Ca_v_1.2α directly as part of a large RNC•MPT•EMC complex. We wanted to provide further proof that such a complex really exists using two complementary approaches. First, we subjected ALFA-eluted stalled Ca_v_1.2α RNCs to sucrose gradient centrifugation to separate ribosome-free from ribosome-bound nascent chains and analyzed co-migration of biogenesis factors with RNCs by western blotting. Indeed, we found that both EMC and Ca_v_β co-migrated with stalled RNCs in the ribosome fraction (**Fig. S7B**). Second, we performed a two-step IP of Ca_v_1.2α 2-TM and 12-TM stall constructs via the ALFA-tag, followed by anti-EMC NbE2 IP to interrogate if ribosomes, MPT and EMC form a single complex (**Fig. 5E, Fig. S7C**). This sequential IP provided additional confirmation that the EMC associates with Ca_v_1.2α-synthesizing ribosome•MPT complexes. The 2-step IP of the 12-TM stall construct further revealed that the EMC-IP step selectively enriched the full-length, two-bundle product, as shorter truncation fragments of the nascent chain predominantly remained in the unbound fraction. These data collectively support a model in which the EMC co-translationally engages Ca_v_1.2α’s N-terminal TM bundle as it is released from the MPT’s lipid-filled cavity (**Fig. 6**).

**Figure 6.**
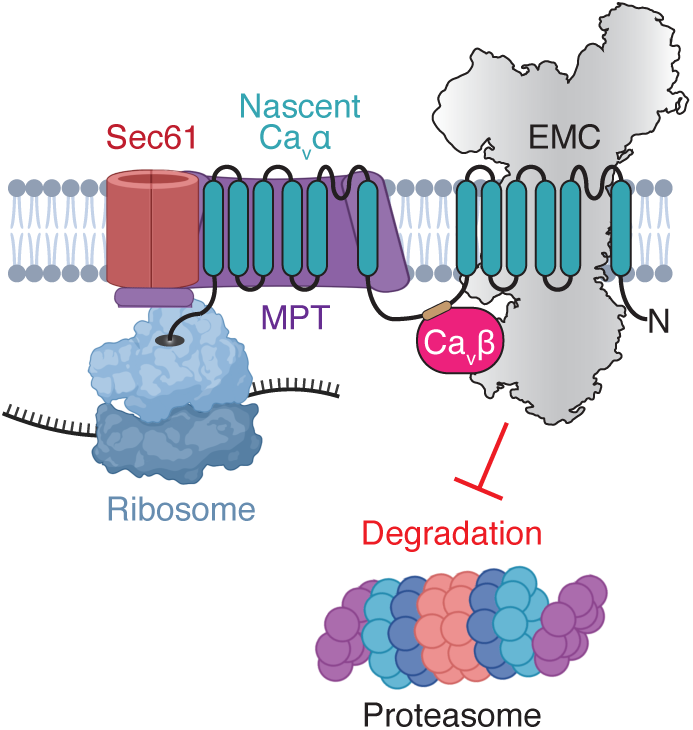
Model for EMC’s chaperone role in calcium channel assembly. The EMC engages nascent calcium channel α-subunits co-translationally to protect the exposed TM bundle I folding intermediate from recognition by quality control factors and so prevents their wasteful premature degradation. This promotes α-subunit folding and co-translational assembly with a matching β-subunit.

## DISCUSSION

Focusing on the cardiac voltage-gated calcium channel Ca_v_1.2, a confirmed direct interactor of the EMC (Chen et al., 2023), we have isolated two separable functions of the EMC in Ca_v_ channel biogenesis. The EMC is required both as an insertase and as a chaperone to protect the nascent Ca_v_1.2α-subunit from premature degradation and so enables its assembly into a functional heterotrimeric channel. We established mutations and nanobodies that selectively inhibit EMC’s chaperone, but not insertase function. Using these tools, we show that EMC’s intramembrane and cytosolic interaction interfaces with the Ca_v_1.2α•β assembly intermediate are required for its chaperone function. Blocking EMC’s chaperone function with two distinct, highly specific inhibitory nanobodies strongly impaired contraction of cardiomyocytes, which endogenously express Ca_v_1.2, providing first evidence for the physiological relevance of EMC’s role as a membrane protein chaperone.

A role for the EMC beyond TM insertion has long been speculated based on the following observations: i) Purification of the EMC from yeast and human cells followed by mass spectrometry revealed multiple interaction partners, some of which could potentially be clients or chaperones, which led to a model in which the EMC functions as a hub for membrane protein biogenesis (Jonikas et al., 2009; Shurtleff et al., 2018; Miller-Vedam et al., 2020). ii) Three proteomic studies showed that many EMC-dependent membrane proteins lack canonical insertase client features and contain multiple TMs, including ion channels and transporters (Shurtleff et al., 2018; Tian et al., 2019; Volkmar et al., 2019). iii) Mutations outside of EMC’s insertase side were shown to impair biogenesis of the N_cyt_ topology membrane protein TMEM97 (Miller-Vedam et al., 2020). iv) Klose *et al*. mapped interactors of the intramembrane chaperone site of the EMC using photocrosslinkers placed in the TM of EMC1 (Klose et al., 2025). Our work is consistent with these previous observations and directly supports a chaperone role of the EMC in calcium channel assembly as suggested by Chen et al. (Chen et al., 2023). Collectively, these findings strongly suggest that EMC’s chaperone function extends beyond Ca_v_ channels to serve other ion channels or classes of membrane proteins.

A more comprehensive client list is needed to help identify common recognition motifs or structural features that mediate dependence on EMC’s chaperone function. Since such interactions might be transient and escape identification using classical affinity purification mass spectrometry approaches, proximity labeling or crosslinking methods might be better suited to capture novel chaperone clients of the EMC (Klose et al., 2025). Any such strategy should be carefully designed to only map interactors of fully assembled EMC complexes rather than overexpressed EMC subunits. Follow-up characterization to validate putative clients should i) establish a direct interaction with the EMC, ii) show that selective perturbation of binding results in client destabilization and iii) control that EMC’s insertase function remains unaffected. Since simple EMC8 depletion is not fully compensated by EMC9, reduces total EMC levels, and thus also compromises EMC’s insertase activity, more targeted and acute approaches will be needed to uncover novel EMC8-dependent chaperone clients. The validated function-separating mutations and inhibitory nanobodies described in this study will greatly facilitate such endeavors in the future. Efficient degradation of a chaperone client in the absence of EMC engagement would, however, require that no other membrane protein chaperone can compensate loss of EMC chaperone function. Although a few other membrane protein chaperones have recently been identified (Gu et al., 2016; Li et al., 2017; Chitwood and Hegde, 2020; Hooda et al., 2024), it is likely that many more still await discovery.

Surprisingly, we found that selective loss of EMC’s insertase function, either directly or indirectly impaired Ca_v_1.2α insertion into the ER membrane. Given the N_cyt_ and C_cyt_ topology of Ca_v_1.2α this was unexpected. Yet, structurally related insertases of the Oxa1 superfamily (Anghel et al., 2017) like Oxa1 and the GEL complex subunit TMCO1 have been proposed to insert internal TMs with such topology in the form of TM hairpins (Chitwood and Hegde, 2019). Further work will be required to establish if purified, reconstituted EMC can indeed insert TM hairpins.

Our data further reveal that EMC complexes containing either EMC8 or its paralog EMC9 are not functionally equivalent and suggest that the vertebrate-specific duplication of the EMC8 gene (Wideman, 2015) allowed the formation of specialized EMC complexes. Only EMC8-containing EMC complexes support calcium channel assembly, since only EMC8 can bind Ca_v_β-subunits. The corresponding interface residues are not preserved in EMC9 (**Fig. S2A**). Interestingly, cell lines of different tissue origins express remarkably different ratios of both paralogs (Uhlén et al., 2015) and EMC9 loss causes distinct developmental defects in both frogs and humans, indicating that EMC8 cannot fully compensate its absence (Jin et al., 2017; Marquez et al., 2023). We therefore propose that EMC9-containing EMC complexes might serve distinct chaperone clients or possess an entirely different function. A careful systematic analysis in cell lines of different tissue origin will be required to determine if EMC complexes containing either EMC8 or EMC9 interact with unique membrane proteins.

Beyond their use to identify novel EMC chaperone clients, we anticipate that the anti-EMC nanobody toolbox will be broadly useful for the membrane protein biogenesis field, as well as labs studying EMC’s role as a viral host factor (Lin et al., 2019; Ngo et al., 2019). The role of EMC’s chaperone function in viral membrane protein biogenesis constitutes an important future research direction.

Our study further revealed that the EMC acts as a chaperone by engaging nascent Ca_v_1.2α as soon as sufficient TMs have emerged from the MPT’s lipid-filled cavity. These findings are consistent with previous observations of EMC•BOS complex interactions (Page et al., 2024) and establish a functional context for collaboration between these large membrane protein biogenesis factors. The lipid-filled cavity of the MPT is formed by the auxiliary GEL, BOS and PAT subcomplexes at the back of the Sec61 complex (McGilvray et al., 2020; Smalinskaitė and Hegde, 2023). The GEL complex contains the EMC3•EMC6 structural homolog TMCO1•OPTI and was proposed to function as the major insertase for internal TM helices of multipass membrane proteins. Using single TM bundle model proteins, the PAT complex was shown to bind inserted TMs with exposed hydrophilic residues and facilitate their folding into TM bundles, which triggered their release from PAT, likely resulting in client dissociation from the lipid-filled cavity (Chitwood and Hegde, 2020; Smalinskaitė et al., 2022). Based on this model and the finite dimensions of the lipid-filled cavity, we postulate that it can only accommodate one TM bundle at a time. For multi-TM bundle proteins this would mean that an upstream TM bundle would need to be released into the lipid bilayer to integrate and fold downstream TM bundles. Ca_v_α-subunits contain four 6-TM bundles that all pack against each other in the final folded channel structure in an intricate domain-swapped architecture. The released α-subunit TM bundle I thus represents a folding intermediate with exposed interface residues that is likely vulnerable to non-specific interactions, aggregation, or recognition as a misfolded quality control substrate. To prevent wasteful premature degradation, TM bundles exposed from the MPT’s lipid-filled cavity might thus require protection against such detrimental fates by co-translational engagement of a membrane protein chaperone, as suggested recently (Sundaram et al., 2025). Our data for Ca_v_1.2α suggest that the EMC can fulfill such a protective role and stabilize nascent multipass membrane protein folding intermediates using its chaperone function. Our data thus extend previous observations of co-translational engagement of multipass membrane proteins by the EMC using ribosome profiling (Shurtleff et al., 2018). We speculate that other yet unidentified chaperones might fulfill similar roles for other classes of multi-bundle membrane proteins.

Steric constraints imposed by EMC’s large cytosolic domain likely prevent the EMC from accessing nascent chains directly at Sec61’s lateral gate, since it would clash with ribosomes at the Sec61•ribosome junction (O’Donnell et al., 2020). In the context of the MPT, however, nascent TM bundles emerge from the lipid-filled cavity instead, whose opening to the lipid bilayer is sufficiently far away to potentially accommodate a co-translationally engaging EMC complex. An important future goal will be to determine how exactly the EMC is recruited to MPT•RNCs complexes.

## ACKNOWLEDGMENTS

CS and GDV were supported by the National Institute Of General Medical Sciences of the National Institutes of Health under Award Number T32GM136631. Cell sorting and flow cytometry analysis for this project was done on instruments in the Stanford Shared FACS Facility (SSFF) (RRID: SCR_017788). We would like to thank SSFF staff members Bianca Gomez and director Lisa Nichols, as well as staff members of Stanford University’s Cryo-electron Microscopy Center (cEMc) and Stanford-SLAC Cryo-EM Facilities (S2C2) for training and troubleshooting advice. We would like to thank Ron Kopito, Rebecca Voorhees, Onn Brandman and Mandi Ma for thoughtful discussions, as well Stanford’s MCP graduate students Mia Greeson, Kyle Trinh and Samantha Johnson, who rotated in the lab and helped with various aspects of this project. Finally, we would like to thank our lab mascot and head of nanobody production, our alpaca Nutmeg, for providing access to its exquisite nanobody immune repertoire. Figure panels 1A, 1F, 2A, 3A, 4B, 5A-B, 6, and S1F-G were created with BioRender.com.

## AUTHOR CONTRIBUTIONS

MS, MB, BS, MA, CES, GDV, AG and TP performed all experiments and data analysis. TP conceived the study. TP wrote the manuscript with input from all authors.

## MATERIALS & METHODS

### Plasmids

Constructs for *in vitro* translations in rabbit reticulocyte lysate were based on the pSP64 vector (Promega, USA). pSpCas9(BB)-2A-Puro (PX459) was a gift from Feng Zhang (Addgene plasmid #48139) (Ran et al., 2013). pLG1-puro non-targeting sgRNA 3 was used as a negative control CRISPRi sgRNA and was a gift from Jacob Corn (Addgene plasmid #109003) (Liang et al., 2018). pJR103 was used to clone dual EMC3 guide RNAs and was a gift from Marco Jost & Jonathan Weissman (Addgene plasmid #187242) (Replogle et al., 2022). The 2nd generation lentiviral packaging plasmid psPAX2 (Addgene plasmid #12260) and envelope plasmid pMD2.G (Addgene plasmid #12259) were gifts from Didier Trono. The pHAGE2 lentiviral transfer plasmid was a gift of Magnus A. Hoffmann and Pamela Bjorkman. The expression plasmid for the SENP^EuB^ protease (Addgene plasmid #149333) was a gift of Dirk Görlich (Vera Rodriguez et al., 2019). Note that the mCherry variant of RFP was used throughout this study, but the simpler nomenclature of RFP is used in the text and figures. Similarly, EGFP is used throughout this study, but referred to as GFP. Superfolder GFP is referred to as sfGFP (Pédelacq et al., 2006). The coding sequence for rat CACNA1C (Ca_v_1.2α) was a gift from Jian Yang (Columbia University, New York, USA). The coding sequences for human CACNB1a, CACNB2b, CACNB3 and rabbit CACNA2D1 were synthesized by Twist Bioscience (USA). The coding sequence for mouse Ca_v_2.1 (CACNA1A) derived from Addgene plasmid #26578, which was a gift from Diane Lipscombe (Richards et al., 2007). The following cDNA clones were purchased from Horizon Discovery (UK): human SOAT1 (clone BC028940), human Cav1.1(CACNA1S) (clone BC133671.1) and human CACNB4 (clone BC075049).

### Antibodies

The following antibodies were used in this study: rabbit polyclonal anti-EMC1 (26017-1-AP, Proteintech, USA); rabbit polyclonal anti-EMC2 (25443-1-AP, Proteintech, USA); mouse polyclonal anti-EMC3 (67205-1-Ig, Proteintech, USA); rabbit polyclonal anti-EMC5 (A305-833, Fortis Life Sciences, USA); rabbit polyclonal anti-EMC8 (19889-1-AP, Proteintech, USA); rabbit polyclonal anti-EMC9 (23919-1-AP; Proteintech, USA); rabbit polyclonal anti-CACNA1C (#ACC-003, Alomone labs, Israel); rabbit polyclonal anti-CACNB3 (#PA5-109280, Thermo Fisher, USA); rabbit polyclonal anti-CACNA2D1 (27453-1-AP, Proteintech, USA); mouse monoclonal anti-α-Tubulin (#T9026, Millipore-Sigma, USA); rabbit polyclonal anti-RPL17 (AP9892b, Abcepta, USA); rabbit polyclonal anti-NOMO2 (14328-1-AP, Proteintech, USA); rabbit polyclonal anti-Ubiquitin (10201-2-AP, Proteintech, USA); mouse monoclonal anti-FLAG M2-HRP (A8592, Millipore-Sigma, USA). The rabbit polyclonal antibody against Sec61β was a gift from Ramanujan Hegde. Secondary antibodies used for western blotting were: LI-COR IRDye 800CW Goat anti-Rabbit IgG Secondary Antibody (NC9401842, Fisher Scientific, USA) and LI-COR IRDye 680RD Goat anti-Mouse IgG Secondary Antibody (NC0252290, Fisher Scientific, USA).

IRDye 680RD-coupled anti-ALFA tag nanobody was generated by expressing the anti-ALFA nanobody Super-Tight variant (Götzke et al., 2019) with a C-terminal Cysteine (Pleiner et al., 2015) and coupling it to IRDye 680RD maleimide (LICOR Biotech, USA) as previously described (Pleiner et al., 2018).

### sgRNAs and siRNAs

The following sgRNAs were cloned into pJR103 for dual guide CRISPRi knockdown of EMC3: EMC3-1 (GCAGTCGCAGGAGAGTTCTG) and EMC3-2 (GAAGCTCGGCTCACAGTCGC).

The following sgRNAs were cloned into pX459 and used to create clonal HEK293 EMC8 KO and HEK293 EMC9 KO cell lines, as well as a stable HEK293 Ca_v_1.2α•Ca_v_β_3_ reporter cell line with EMC5 KO: EMC5 (GCATCATGGCGCCGTCGCTG), EMC8 (GTGGCCTCCAGAATCGCCGA) and EMC9 (CAAAAACAGCCCGTTGACTG).

The following siRNAs were used in this study: negative control no. 2 siRNA (#4390846), EMC1 s225925, EMC8 siRNA s20201, EMC9 siRNA s27245 (Silencer Select; Thermo Fisher Scientific, USA).

### Expression and purification of proteins

GFP- and ALFA-tagged proteins were purified from human cell lysate using protease-cleavable biotinylated anti-GFP and anti-ALFA tag nanobodies (Kirchhofer et al., 2010; Götzke et al., 2019). These nanobodies were expressed in *E. coli* and purified using Ni^2+^-chelate affinity chromatography as described in detail before (Pleiner et al., 2015; Pleiner et al., 2020; Stevens et al., 2024).

Immobilized biotinylated nanobodies were eluted from streptavidin magnetic beads (#88817, ThermoFisher Scientific, USA) using an engineered SUMO protease (SENP^EuB^) that recognizes the SUMO^Eu1^ module (Vera Rodriguez et al., 2019) or SUMOstar protease that recognizes the SUMOstar module (Liu et al., 2008). Both proteases are orthogonal. His_14_-Tev-tagged SENP^EuB^ protease (Addgene ID #149333) was expressed in *E. coli* NEB express I^q^ as described before (Stevens et al., 2024).

His_14_-*bd*NEDD8-tagged human Ca_v_β_3_ was expressed for 6h at 18°C in *E. coli* Rosetta-gami 2 with 0.2 mM IPTG induction. The protein was purified using Ni^2+^-chelate affinity chromatography. The *bd*NEDD8 tag was cleaved right before use of the protein in *in vitro* translation reactions by incubation with 300 nM of the cognate *bd*NEDP1 protease for 20 min. at 4°C (Frey and Görlich, 2014).

### Mammalian *in vitro* translation

*In vitro* translation reactions in rabbit reticulocyte lysate (RRL) were carried out with *in vitro* transcribed mRNA as described before (Sharma et al., 2010). PCR products generated from pSP64-derived plasmids or gene fragments (Twist Biosciences, USA) served as templates for run-off transcription and contained a 5’ SP6 promoter followed by an open-reading frame and a 3’ stop codon. A 10 µl transcription reaction contained 7.6 µl T1 mix (Sharma et al., 2010), 0.2 µl SP6 polymerase (New England Biolabs, USA), 0.2 µl RNAsin (Promega, USA), 100 ng PCR product, and was carried out for 1.5h at 37°C. Transcriptions were added directly to homemade RRL at 5% v/v. RRL was treated with micrococcal nuclease S7 (#10107921001, Roche, Germany) in the presence of CaCl_2_ to remove endogenous hemoglobin mRNA and then inactivated with EGTA. Nascent proteins were labeled during translation reactions of ∼10 min/10 kDa at 32°C in RRL by incorporation of radioactive ^35^S-methionine (Revvity, USA). To enable ER membrane incorporation of translated membrane proteins, RRL is supplemented with 5% (v/v) of human ER-derived microsomes (hRMs), prepared as described below. Samples were analyzed by SDS-PAGE and autoradiography to detect the translated ^35^S-labeled proteins.

### Protease protection assay

To assess the membrane spanning topology of Ca_v_1.2α TM bundle 1, a PCR product comprising amino acids 79-503 and ending in 3x stop codons was generated as template for *in vitro* transcription. The channel further contained an N-terminal 3xFLAG tag as well as an internal 1xHA tag that was inserted in the lumenal loop between TM1 and TM2 at position E149 (Uniprot: P22002-5). This construct was then translated in RRL in the presence of HEK293 wild-type or EMC5 KO hRMs (Pleiner et al., 2023) as described above. Protease-accessible regions were digested by incubation with 0.5 mg/ml Proteinase K for 50 min. at 4°C in the presence or absence of 0.05% (v/v) Triton-X-100 to solubilize hRM membranes. Proteinase K was inactivated by addition of 5 mM PMSF and quick transfer into boiling SDS buffer (100 mM Tris/HCl pH 8.4; 1% [w/v] SDS). Denatured digestion reactions were diluted tenfold with IP buffer (50 mM HEPES/KOH pH 7.5; 300 mM NaCl; 0.5 % [v/v] Triton-X-100) and incubated with anti-HA or anti-FLAG M2 resin (Millipore-Sigma, USA) for 1 hour at 4°C for immunoprecipitation of protected fragments. After washing with IP buffer, bound fragments were eluted with SDS-PAGE sample buffer.

### Preparation of human ER-derived microsomes (hRMs)

To prepare hRMs from Expi293, HEK293 WT or EMC5 KO cell lines, we followed a recently described procedure (Sundaram et al., 2022). Briefly, cells were harvested and then washed twice in 1x PBS. Cells were then resuspended in 4x pellet volume of hypotonic buffer (10 mM HEPES/KOH pH 7.5.; 10 mM KAc, 1 mM MgAc; 1x Protease inhibitor cocktail [Roche, Germany]) and lysed with 6 passages through a cell cracker using a 16 µm-clearance ball bearing (Isobiotec, Germany). Complete cell lysis was verified by trypan blue staining. The lysate was then supplemented with 250 mM sucrose and spun twice for 3 min at 1500 g in a table-top centrifuge (5430R, Eppendorf, Germany) at 4°C to remove nuclei and cell debris. The resulting supernatant was then centrifuged for 10 min. at 10,000 g at 4°C in a table-top centrifuge (5430R, Eppendorf, Germany). The supernatant was aspirated, and the membrane pellet gently resuspended in nuclease buffer (10 mM HEPES/KOH pH 7.5; 250 mM KAc, 10 mM MgAc; 250 mM Sucrose). To remove endogenous mRNAs, membranes were further treated with 4U/µL Micrococcal nuclease (#M0247S, New England Biolabs, USA) in the presence of 1 mM CaCl_2_, 2 U/mL TurboDNAse (#AM2238, Thermo Fisher Scientific, USA) and 0.5 mM PMSF for 10 min in a 37°C water bath, and then quenched by Ca^2+^-chelation with 2 mM EGTA for 2 min. at 23°C. Nucleased membranes were pelleted for 10 min. at 10,000 g at 4°C as above and resuspended in microsome buffer (20 mM HEPES/KOH pH 7.5; 280 mM sucrose, 0.5 mM EDTA; 2 mM DTT, 1x Protease inhibitor cocktail) containing 40 U/mL Superase-In (#AM2694, Thermo Fisher Scientific, USA). After a 5 min. incubation at 23°C, membranes were pelleted as above and resuspended in microsome buffer. The absorbance at 280 nm of the resuspended membranes was measured by boiling an aliquot in SDS buffer. The hRM preparation was then adjusted to an absorbance of 50 at 280 nm using microsome buffer. Nucleased hRMs were used fresh or snap-frozen in liquid nitrogen in single-use aliquots and stored until further use at −80°C.

### Purification of stalled ribosome nascent chain complexes

To generate stalled, ribosome-attached nascent chains of defined length, PCR products for run-off transcriptions lacked stop codons and encoded a C-terminal MLKV appendage as described before (Smalinskaitė et al., 2022). The following construct boundaries were chosen to generate Ca_v_1.2α biogenesis intermediates stalled such that a defined number of TMs were ribosome exposed, accounting for ∼35 amino acids (aa) hidden in the ribosome exit tunnel: 2xTMs (aa79-219+MLKV); 4xTMs (aa79-292+MLKV); 6xTMs (aa79-442+MLKV); 6xTMs+AID (aa79-537+MLKV); 12xTMs (aa79-786+MLKV). All constructs are based on the rat Ca_v_1.2α channel (Uniprot: P22002-5) and additionally carried a N328A mutation to abolish a glycosylation acceptor site, as well as an N-terminal ALFA tag. The ASGR1 single-pass stalled control construct is based on human ASGR1 and comprised aa1-147+MLKV and also contained a N79A glycosylation site mutation and an N-terminal ALFA tag.

*In vitro* transcription and translation were performed as described above. ∼500 µL translation reactions were supplemented with 5% (v/v) hRMs and 150 nM purified, cleaved *hs*Ca_v_β_3_. After translation, hRMs were pelleted through a 20% (w/v) sucrose cushion prepared in 1x Physiological salt buffer (1xPSB) (25 mM HEPES/KOH pH7.5, 100 mM KAc, 5 mM MgAc) by spinning for 10 min. at 12,500 g at 4°C in a table-top Eppendorf 5430R centrifuge (Eppendorf, Germany). The supernatant was aspirated and the hRM pellet resuspended in 500 µL 1x PSB. Resuspended hRMs were then diluted with 500 µL solubilization buffer (1xPSB, 3.5% w/v Digitonin, 1x Protease inhibitor cocktail [Roche, Germany]) and incubated for 30 min. rotating head-over-tail at 4°C. Insoluble material was spun out for 10 min. at 16,000 g at 4°C. The supernatant was then added to magnetic Streptavidin beads containing immobilized, biotinylated SUMOstar-cleavable anti-ALFA nanobody (Stevens et al., 2024) that were additionally equilibrated in wash buffer (50 mM HEPES/KOH pH7.5, 150 mM NaCl, 10 mM MgAc, 0.25% w/v Digitonin, 1 mM DTT) and blocked with excess biotin. After 1h binding at 4°C, beads were washed three times with 500 µL wash buffer and ALFA nanobody-bound RNCs were eluted by incubation with 500 nM SUMOstar protease in wash buffer at 4°C for 30 min. The eluate was then layered over 300 µL sucrose cushion buffer (500 mM sucrose in wash buffer) and spun for 1h at 100,000g at 4°C in a TLA120.1 rotor to isolate the ribosome-associated fraction. The ribosome pellet was resuspended in 30 µL sucrose cushion buffer. The ribosomal rRNA content of the different samples was then normalized by A_260_ absorption. Normalized samples were analyzed by western blotting.

### Sucrose gradient centrifugation

Stalled ALFA-tagged Ca_v_1.2α biogenesis intermediates were generated and purified with anti-ALFA nanobody as described above. 200 µL protease elution were then layered on top of a 2 mL 10–50% (w/v) sucrose gradient prepared in wash buffer and centrifuged for 1 hour at 55,000 rpm at 4 °C in a TLS-55 rotor (Beckman Coulter, USA) using the lowest acceleration and deceleration settings. 11x 200 µl fractions were harvested from the top of the gradient and samples of each fraction were analyzed by western blotting.

### Cell culture

Adherent HEK293 cell lines and Lenti-X cells were cultured in Dulbecco’s Modified Eagle Medium (DMEM) supplemented with 10% fetal bovine serum (FBS) and 2 mM L-Glutamine. hTERT RPE1 dCas9-BFP-KRAB CRISPRi cells (Jost et al., 2017) were cultured in DMEM/F-12 (1:1) supplemented with 10% FBS and 2 mM L-Glutamine and used throughout this study. For simplicity, we simply refer to this cell line as RPE1 cells. Expi293 cells (Thermo Fisher Scientific, USA) were maintained at a concentration of 0.5-2.0 million cells per ml in Expi293 Expression Medium (Thermo Fisher Scientific, USA).

The cardiomyocyte differentiation protocol was adapted from Lian et al., 2013 (Lian et al., 2013). The human iPSC line expressing the fluorescent calcium indicator GCaMP6f used for our experiments was described previously (Huebsch et al., 2015). The cell line was maintained in mTeSR Plus (STEMCELL Technologies, Canada; cat. no. 100-0276) and passaged with Accutase at ∼70% confluency. For differentiation, cells were seeded into 12-well plates at a density of 100,000 cells per well in mTeSR Plus supplemented with ROCK inhibitor (Tocris Bioscience, United Kingdom, cat. no. 1254). Cells were allowed to recover overnight. On day 0, the medium was exchanged to RPMI (Thermo Fisher Scientific, USA, cat. no. 11875119), B-27 minus insulin (Thermo Fisher Scientific, USA, cat. no. A1895601) supplemented with 6 μM CHIR99021 (Tocris Bioscience, United Kingdom, cat. no. 4423). After 48 hours of CHIR exposure, the medium was exchanged to RPMI containing 5 μM IWP4 (Tocris Bioscience, United Kingdom, cat. no. 5214) and refreshed every 24 hours.

On day 5, differentiating cultures were reseeded at a 1:1 ratio, and immediately transduced with lentivirus encoding the specific nanobody constructs in presence of 6 µg/mL polybrene (Sigma-Aldrich, USA, cat. no. TR-1003). The medium was exchanged 48 hours after transduction and subsequently refreshed every 24 hours.

### CRISPRi knockdowns

RPE1 dCas9-BFP-KRAB wild-type or stable RPE1 dCas9-BFP-KRAB cells expressing Ca_v_1.2-GFP-2A-RFP and Ca_v_b_3_-miRFP713 were transduced with lentivirus containing a pLG1-puro non-targeting sgRNA 3 or pJR103 backbone encoding dual anti-EMC3 sgRNAs. Sequences of sgRNAs were derived from the hCRISPRi-v2 compact library (Horlbeck et al., 2016). 48 h after transduction, 1 µg/ml puromycin was added for three consecutive days to select cells with a successfully integrated sgRNA expression cassette. After two days of recovery, cells were transduced with fluorescent reporters expressed from a lentiviral backbone under control of a CMV promoter. Cells were analyzed 48 h after reporter transduction by flow cytometry (8 days after sgRNA transduction).

### Lentiviral transduction

Lentivirus was generated by co-transfection of Lenti-X cells (Takara Bio, Japan) with a desired transfer plasmid and two packaging plasmids (psPAX2 and pMD2.G) using the TransIT-293 transfection reagent (Mirus, USA). 48 h post transfection, culture supernatant was harvested, aliquoted and flash frozen in liquid nitrogen. For lentiviral transduction of adherent RPE1 cells, 50-200 µl lentiviral supernatant and 8 µg/ml polybrene (Millipore-Sigma, USA) were usually added directly to ∼70% confluent cells in 2.5 ml culture medium in a 6-well.

All lentiviral fluorescent reporter constructs were generated in the pHAGE2 backbone and expressed from a CMV promoter. The tail-anchor model clients human FDFT1/SQS (EMC client) and human VAMP2 (GET1/2 client) contained their TMD and flanking regions fused to the C-terminus of RFP in a GFP-2A-RFP cassette as described before (Guna et al., 2018; Pleiner et al., 2020). Multipass EMC insertase client reporters were generated by fusing the complete coding sequence of human GPCR AGTR2 (N_exo_ client) to the N-terminus of EGFP in a GFP-2A-RFP cassette, as well as the complete coding sequence of human SOAT1 (C_exo_ client) to the C-terminus of GFP in the same cassette. Sec61-dependent ASGR1 control reporter was generated by fusing the coding sequence of human ASGR1 to the C-terminus of RFP in our GFP-2A-RFP cassette as described before (Chitwood et al., 2018; Pleiner et al., 2023).

EMC3 WT or EMC3 Mut add-back constructs were expressed from a EMC3-ALFA-2A-TagBFP-3xFLAG cassette driven by a weak PGK promoter. EMC8 WT, EMC8 Mut or EMC9 were expressed from a 3xFLAG-TagBFP-2A-ALFA-(GlySer)_11_-EMC8 cassette driven by a weak PGK promoter. EMC1 WT and EMC1 Mut lacking their original signal sequence (23-end) were expressed from a 3xFLAG-TagBFP-2A-Prl(ss)-ALFA-EMC1(23-end) cassette (ALFA = ALFA peptide tag, Prl = Prolactin, ss = signal sequence) driven by a weak PGK promoter. Nanobody expression cassettes for cytosolic expression comprised a 3xFLAG-TagBFP2-2A-ALFA-nanobody open reading frame and were driven by a CMV promoter. To secrete nanobodies across the ER membrane and retain them in the ER lumen we used a 3xFLAG-TagBFP2-2A-Prl(ss)-ALFA-nanobody-KDEL cassette (KDEL = Lys-Asp-Glu-Leu ER retention signal) driven by a CMV promoter.

Lentiviral constructs for cardiomyocyte transduction with nanobodies contained an upstream ubiquitous chromatin opening element (UCOE) followed by an EF1α promoter. The open reading frame encodes TAGBFP2 separated by a viral 2A site from the respective control or inhibitory anti-EMC nanobodies. Transduced cardiomyocytes are thus marked by BFP expression.

### Stable cell line generation

To study calcium channel stability in human cells the following stable cell lines were generated by lentiviral transduction of RPE1 dCas9 CRISPRi cells and cell sorting via the appropriate fluorescent protein markers: *rn*Cav1.2α(ΔC,aa1-1662)-GFP-2A-RFP alone, or co-expressed with either *hs*CACNB3(fl)-miRFP713 or *hs*CACNB3(fl)-miRFP713 fused via a P2A sequence (de Felipe et al., 2006) to Prl(ss)-3xHA-*oc*CACNA2D1(aa29-end). The C-terminal truncation of Ca_v_1.2α distal to the Ile-Gln (IQ) domain was previously characterized to assemble into fully functional heterotrimeric calcium channel complexes and possesses properties identical to the full length α-subunit as shown by electrophysiology (Chen et al., 2023). Similar C-terminal deletions improved expression levels of human CACNA1S, thus the reporter contained *hs*Ca_v_1.1α (ΔC,aa1-1434)-GFP-2A-RFP, and of mouse CACNA1A, with the reporter containing *mm*Ca_v_2.1α(aa88-1934)-GFP-2A-RFP, additionally avoiding an unstructured N-terminal region with extremely high GC content that prevented PCR amplification. Both *hs*Ca_v_1.1 and *mm*Ca_v_2.1 were co-expressed with *hs*CACNB3(fl)-miRFP713. Polyclonal cell populations expressing all desired reporters were sorted using a SH800S cell sorter (Sony Biotechnology, Japan).

### siRNA rescue assays

72 hour siRNA knockdowns were performed by reverse transfection of either 150,000 RPE1 cells or 300,000 HEK293 cells in 6-well plates with 10 nM siRNA complexes formed in Opti-MEM Reduced Serum Medium using RNAiMax transfection reagent (Thermo Fisher Scientific, USA). 24h post-transfection the medium was exchanged and ∼50-500 µL lentiviral supernatant for reporter and/or rescue constructs was added to the 6-well together with 8 µg/mL polybrene. 24h later cells were split 1:2 if necessary.

### Confocal Microscopy

BFP-positive (lentivirus-transduced) cardiomyocytes were imaged between days 9–10 using a Zeiss LSM 990 confocal microscope. Live-cell calcium imaging was performed under controlled temperature and CO₂ conditions, using identical exposure, laser power, and frame-rate settings for all samples to allow quantitative comparison of GCaMP fluorescence dynamics.

### Image analysis

Calcium transient analysis was performed on exported video files using Fiji/ImageJ. For each video, BFP-positive (lentivirus-transduced) cardiomyocytes were identified based on their blue fluorescence, and ROIs were drawn around individual cells in ImageJ. Mean fluorescence intensities were extracted across all frames to generate a time-series trace for each cell. Traces were imported into Python for analysis. Three metrics were quantified: (1) percentage of beating cells, defined by the presence of detectable calcium transients; (2) beating rate (peaks per minute), determined by detecting ΔF/F₀ peaks using a noise-adaptive threshold; and (3) calcium transient amplitude, computed as the maximum ΔF/F₀ following baseline normalization (F₀ = 10th-percentile fluorescence). Peak detection, ΔF/F₀ calculations, and plotting were performed using Python scripts and Matplotlib.

### Purification of GFP- or ALFA-tagged proteins from human cells

Cell pellets were resuspended and incubated with ∼7 ml of solubilization buffer (50 mM HEPES/KOH pH 7.5, 200 mM NaCl, 2 mM MgAc, 1% w/v GDN, 1 mM DTT and 1x protease-inhibitor cocktail [Roche, Germany]) per 1 g cell pellet for 30 min under constant agitation at 4°C. Cell lysates were spun for 10 min. at 16,000xg and 4°C in a table-top Eppendorf 5430R centrifuge to remove cell debris. The protein concentration of the supernatant was determined by A280 nm absorption using a Nanodrop spectrophotometer. SDS-PAGE samples of the total cell lysate were taken such that 40 µg of total protein are loaded in 10 µL per gel lane. In parallel, Pierce magnetic Streptavidin beads (Thermo Fisher Scientific, USA) were i) pre-equilibrated with wash buffer (solubilization buffer with 0.01 % [w/v] GDN), ii) incubated with ∼2 µg biotinylated anti-GFP or anti-ALFA nanobody per confluent T175 flask of cells and then iii) empty biotin binding sites were blocked with 100 µM biotin in wash buffer for 5 min. on ice. The cleared supernatant was then added to nanobody containing, blocked magnetic Streptavidin beads and incubated for 1h at 4°C under constant agitation. Beads were retrieved using magnetic racks (Sergi Lab Supplies, USA) and washed 3x with 1 mL wash buffer, resuspended in ∼20 µL wash buffer containing either 250 nM SENP^EuB^ or 500 nM SUMOstar protease and incubated for 20 min. on ice for elution. One fifth of the eluate was typically analyzed by SDS-PAGE and Sypro Ruby staining. The amount of recovered bait in each sample was typically quantified in ImageJ and this quantification was used to normalize elution samples for western blotting.

### Flow cytometry analysis

RPE1 cells transduced with lentiviral reporters were typically analyzed by flow cytometry after 48 h. Cells were trypsinized, washed, and resuspended in 1xPBS for flow cytometry analysis. Analysis was either on a Quanteon or Penteon benchtop analyzer (Agilent Technologies, USA). We typically collected at least 30,000 cells in the final gate. Flow cytometry data was analyzed using FlowJo v10.8 Software (BD Life Sciences, USA). Unstained cells transiently transfected with either BFP, GFP, RFP or miRFP713 if needed were analyzed separately along every run as single-color controls for multicolor compensation using the FlowJo software package.

Day 10 cardiomyocytes were stained at the concentration of 1.5 × 10⁶ cells/mL. Cells were dissociated, pelleted, and resuspended in FACS staining buffer (BD Biosciences, 554657). Cells were incubated on ice for 2 hours with anti–Cav1.2 primary antibody (Alomone Labs, ACC-130), followed by a wash with staining buffer and incubation with the secondary antibody (Abcam, ab150083). After two additional washes, cells were resuspended in PBS and analyzed by flow cytometry.

### Purification of EMC for alpaca immunization

EMC complex was purified in LMNG detergent micelles from a stable Expi293 suspension cell line expressing GFP-3C-EMC2 using a SENP^EuB^-cleavable anti-GFP nanobody as described before (Pleiner et al., 2020; Stevens et al., 2024). The nanobody eluate was cleaved overnight at 4°C with homemade, purified HRV 3C protease to cleave off the GFP tag. Excess GFP and nanobody were removed by size-exclusion chromatography on a Superose 6 Increase 3.2/300 column. Pure fractions containing untagged EMC complexes were pooled and concentrated for alpaca immunization.

In parallel, using the same strategy EMC was purified in Deoxy Big Chap (Calbiochem, USA) and reconstituted into proteoliposomes containing PE:PC:cholesterol as described before (Guna et al., 2018; Guna et al., 2022).

In addition, the soluble domain of the human EMC comprising full length EMC2, EMC8, EMC5 C-terminus (aa66-end), EMC3 coiled-coil (aa35-117), EMC3 C-terminus (aa196-end) and EMC4 N-terminus (aa2-56) were expressed and purified from *E. coli* and mixed to form a stable complex as described before (Pleiner et al., 2021). The complex was purified by size-exclusion chromatography on a Superdex 200 10/300 column.

All three preparations were injected five times every two weeks at separate locations on the shoulder of a naïve, male alpaca, including an additional shot of GERBU P adjuvant. Alpaca husbandry and immunizations were performed by Antibodies Inc. (Davis, USA). Peripheral blood lymphocytes were isolated after the final boost. Total RNA was extracted, reverse transcribed and used to PCR amplify the nanobody immune repertoire and generate a nanobody library for phage display as described before (Pleiner et al., 2015). Since this first round of immunization revealed limited diversity of the nanobody repertoire, we reimmunized the same alpaca after 6 months with two shots of purified full-length EMC in LMNG and proteoliposomes and prepared a second library.

### Selection of anti-EMC nanobodies by phage display

For nanobody selections from purified bacteriophage libraries, we immobilized the EMC freshly from GFP-3C-EMC2 Expi293 cell lysate using anti-GFP nanobody as described above. After washing the beads, the immobilized EMC was incubated with purified bacteriophage nanobody library to enrich those phages presenting anti-EMC nanobodies. Weak or non-specifically binding phages were washed away and EMC-bound phages were eluted using the SENP^EuB^ cleavage site on the GFP nanobody. We performed three selection rounds using decreasing amounts of cell lysate as input for EMC immobilization. This initial selection yielded a limited set of anti-EMC nanobodies, with one nanobody class (containing insert NbE2) dominating the selection outcome. After re-immunization as described above, we blocked this dominant epitope by immobilizing EMC from wild-type Expi293 cell lysate for phage display selection using biotinylated, SENP^EuB^-cleavable NbE2. This resulted in substantially increased nanobody diversity.

Nanobody coding sequences were amplified from bacteriophages after three rounds of selection by a two-step PCR protocol to add sequencing adaptors and barcodes for deep sequencing on a Illumina MiSeq using the MiSeq Reagent Nano Kit v2 (500-cycles). In parallel, nanobodies were also cloned into a ALFA-sfGFP-SUMO^Eu^-Nb-His_10_ *E. coli* expression vector using Gibson assembly and expressed in 96 deep-well plate format for characterization by ELISA. Positive classes were immobilized directly out of *E. coli* cell lysate using biotinylated, SUMOstar-cleavable anti-ALFA nanobody for small-scale test purifications from Expi293 detergent cell lysate to assess affinity and specificity. After washing, anti-EMC nanobody along with bound EMC (if positive) were specifically eluted using SENP^EuB^ cleavage.

### Cryo-EM grid preparation and data collection

4 µl of purified EMC•NbG9•NbE2 complex in GDN detergent at 1.5 mg/ml concentration was applied to freshly glow discharged Quantifoil R1.2/1.3 grids copper 200 mesh grids. The grids were glow discharged using Pelco easiGlow. The grids were quickly blotted for 3 sec for any excess liquid with a blot force of 3 and wait time of 5 sec and subsequently plunge frozen in liquid ethane using Vitrobot Mark IV (Thermo Fisher Scientific, USA). The vitrified sample was stored in long term storage in liquid nitrogen until imaged. The frozen grids for EMC•NbG9•NbE2 complex were screened using a Glacios 200kV microscope equipped with a Falcon 4i detector. The grid with desired ice thickness was used for data collection on Titan Krios G2 operated at 300 kV and equipped with Falcon 4i detector and post column energy filter, SelectrisX. The data were collected with calibrated pixel size of 0.92, defocus range of −0.6 to −2 micron and a total dose of 54.23 e/Å2 at a dose rate of 8.93 e/px/sec.

### Image processing

The dataset of 16392 micrographs was collected in .tiff format and imported to Cryosparc (Punjani et al., 2017) for preprocessing including patch Motion Correction and Patch CTF estimation. 9348 micrographs were selected based on CTF fit cut-off at 5 and total full frame motion distance within 50 pixels. A picking model was trained on the manually picked particles using Topaz Train (Bepler et al., 2019) and particles were extracted from all curated micrographs based on the model. The box size used for particle extraction was 480 pixels. One round of 2D classification was done followed by 2D selection. Clean classes were subjected to two rounds of multiclass ab-initio reconstruction followed by heterogenous refinement. The class representing the EMC complex bound to both nanobodies was selected and a final round of multiclass ab-initio reconstruction was performed to remove junk particles. The best class was used to create a mask and a non-uniform refinement routine was followed. To further improve the map quality, particles were passed through reference-based motion correction and used for another round of non-uniform refinement. DeepEMhancer (Wrapper) (Sanchez-Garcia et al., 2021) was used in cryoSPARC to sharpen the map and guide model building. Local resolution estimation and resolution-based filtering was performed on the final map.

### Model building and refinement

Initial models were based on a previously published cryo-EM structure (PDB: 8S9S; Pleiner et al., 2023) of the EMC complex, as well as predicted structures of individual subunits of the EMC complex and the Nbs G9 and E2 using Alphafold. The complex was fitted to the Non-uniform refined map using ChimeraX and further manually built using Coot (v0.9.8.86) (Emsley et al., 2010) and iteratively refined by Phenix real space refinement (Liebschner et al., 2019). Table S2 provides details of the cryo-EM map and the model refinement. FSC curves and selected map-model agreement panels are provided in Figure S4F.

**Figure S1.**
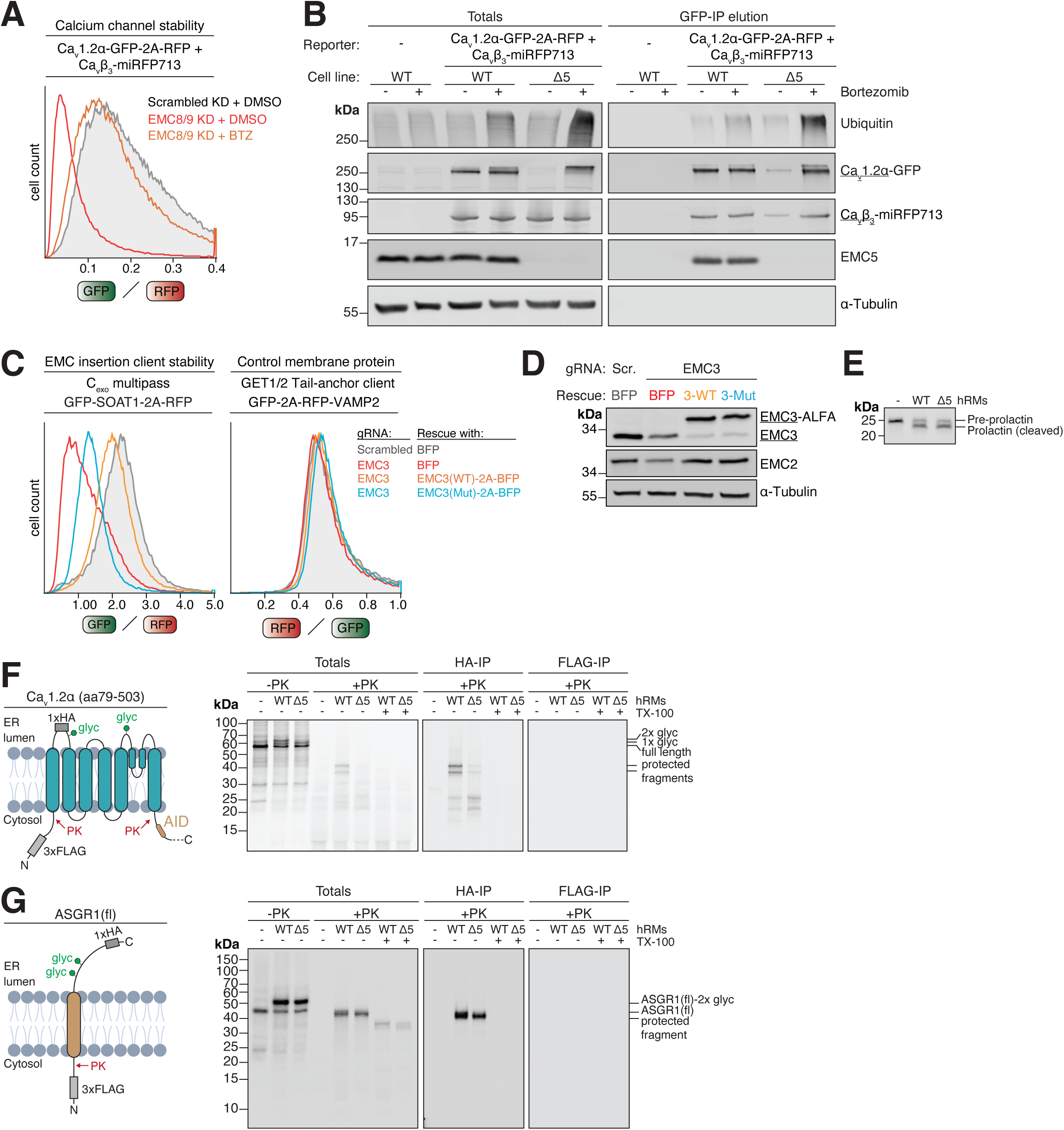
Characterization of Cav1.2α’s EMC dependence. **A)** 72h scrambled or EMC8+9 double siRNA knockdown in stable RPE1 Ca_v_1.2α•Ca_v_β_3_ reporter cells. Cells were additionally treated with 10 nM proteasome inhibitor bortezomib or DMSO solvent control for 16h before analysis of the GFP/RFP ratio by flow cytometry. **B)** ΗΕΚ293 WT, or HEK293 WT and EMC5 KO (Δ5) cells stably expressing the Ca_v_1.2α•Ca_v_β_3_ reporter depicted in Fig. 1B were treated with DMSO or 10 nM bortezomib for 16h. Cells were harvested, lysed in detergent and subjected to anti-GFP nanobody purification of Ca_v_1.2α-GFP. Total cell lysates and protease elution of the GFP-IP were analyzed by western blotting with the indicated antibodies. Note that Ca_v_1.2 α-GFP is largely degraded in EMC5 KO cells, but can be stabilized by bortezomib treatment. The Ca_v_1.2α channels that accumulate are heavily ubiquitinated, indicating that Ca_v_1.2α is ubiquitinated and degraded in the absence of the EMC. **C)** Knockdown of EMC3 by CRISPRi in RPE1 dCas9-BFP-KRAB cells transiently transduced with C_exo_ multipass EMC insertase client reporter SOAT1 (Wu *et al*., 2024) or EMC-independent tail-anchored membrane protein reporter VAMP2, a client of the GET1/2 insertase. Knockdown was rescued with either just BFP or BFP separated by a 2A site from ALFA-tagged wild-type (WT) EMC3 or the insertase-deficient EMC3 R31A+R180A mutant (Mut). The GFP/RFP (SOAT1) or RFP/GFP (VAMP2) ratios of BFP^+^ cells were determined by flow cytometry and are depicted as histograms. **D)** Experiment as in C, but analysis of total cell lysates by western blotting with the indicated antibodies. **E)** WT and EMC5 KO (Δ5) hRMs are equally active in Sec61-dependent protein translocation. ^35^S-methionine-labled bovine preprolactin carrying a cleavable signal sequence was translated in rabbit reticulocyte lysate in the presence or absence of human rough ER membranes (hRMs) prepared from either wild-type or Δ5 HEK 293 cells. **F)** Insertion defect of Ca_v_1.2α in EMC5 KO (Δ5) membranes. As in Fig. 1F, but with Ca_v_1.2α (amino acids [aa] 79-537) carrying both an N-terminal 3xFLAG tag, as well as a 1x HA tag inserted into the lumenal loop between TM1 and TM2. The latter makes an otherwise occluded (Fig. 1F) glycosylation (glyc) site accessible. Non-incorporated as well as cytosolically accessible protein portions were digested with proteinase K (PK) in the presence or absence of Triton-X-100 (TX-100) to solubilize the hRM membrane. The resulting protease protected fragments (PFs) were subjected to denaturing anti-HA and anti-FLAG immunoprecipitations (IP). **G)** No insertion defect of ASGR1, a Sec61-dependent single-pass type II membrane protein, was observed in Δ5 hRMs. Assay as in F, but with full length (fl) ASGR1 carrying an N-terminal 3xFLAG and C-terminal 1xHA tag.

**Figure S2.**
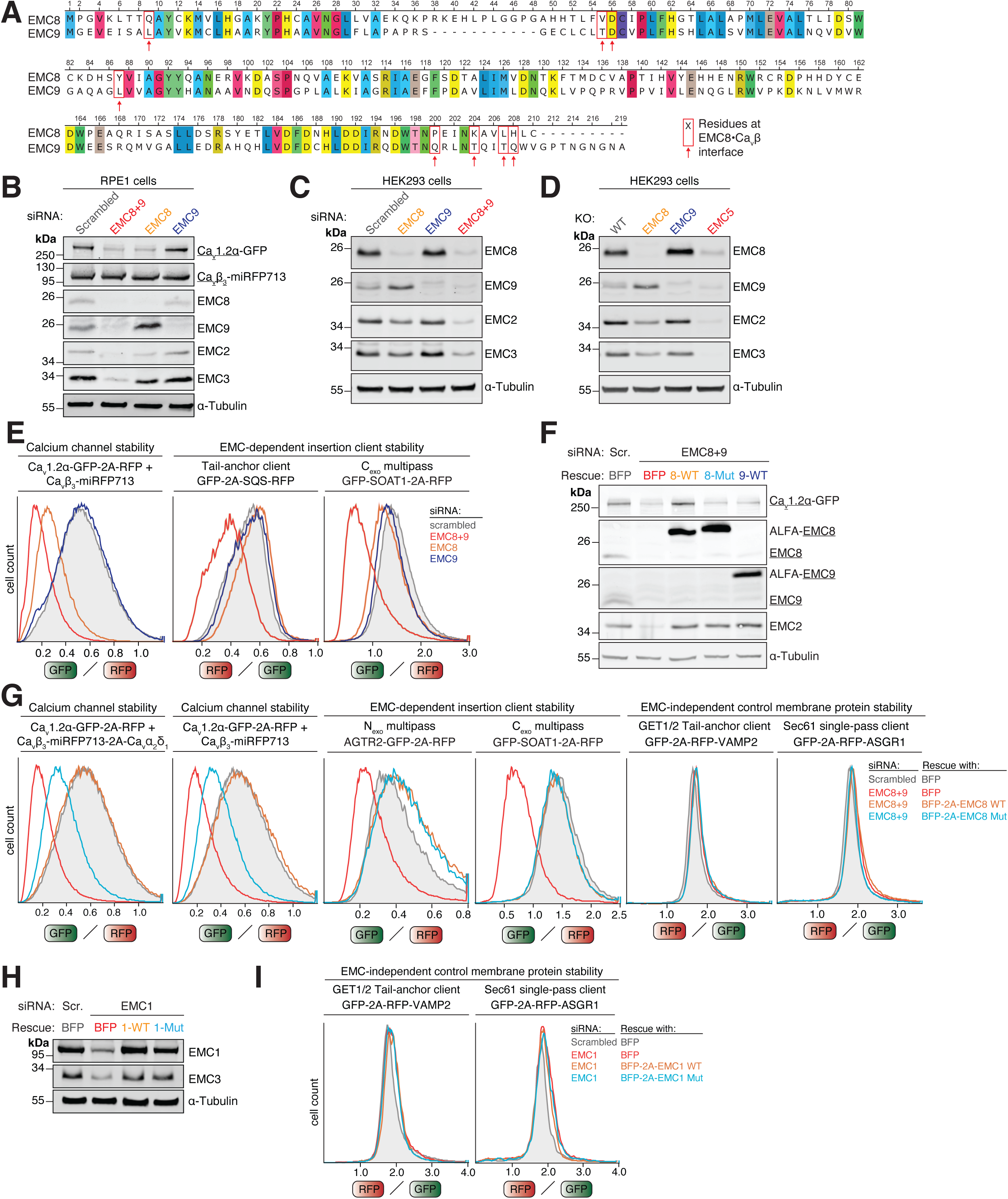
Characterization of EMC’s chaperone interfaces. **A)** Sequence alignment of human EMC8 and EMC9 generated with Clustal Omega (Madeira *et al*., 2024) and visualized with Unipro UGENE (Okonechnikov *et al*., 2012). Both paralogs display ∼40% sequence identity. Identical positions are highlighted. EMC8 residues that contact Ca_v_β_3_ in the EMC•Ca_v_1.2α•Ca_v_β_3_ co-structure (PDB 8EOI, Chen *et al*., 2023) are highlighted in red boxes to illustrate that 7/8 of these residues are altered in EMC9. **B)** 72h scrambled, EMC8, EMC9 or EMC8+9 double siRNA knockdown in RPE1 cells. Total cell lysates were analyzed by western blotting with the indicated antibodies. **C)** As in B, but in HEK 293T cells. **D)** Western blot analysis of total cell lysates prepared from HEK 293T WT, EMC8 KO, EMC9 KO or EMC5 KO cells. **E)** 72h scrambled, EMC8, EMC9 or EMC8+9 double siRNA knockdown in stable RPE1 Ca_v_1.2α•Ca_v_β_3_ reporter cells or RPE1 cells transiently transduced with EMC-dependent insertase client reporters SQS and SOAT1. The GFP/RFP (Ca_v_1.2α, SOAT1) or RFP/GFP (SQS) ratios of the resulting cells were determined by flow cytometry and are depicted as histograms. **F)** 72h scrambled or EMC8+9 double siRNA knockdown in stable RPE1 Ca_v_1.2α•Ca_v_β_3_ reporter cells. 24h post siRNA transfection, cells were transduced with lenti-viral rescue constructs encoding either just BFP or BFP separated by a 2A site from ALFA-tagged EMC8 WT, EMC8 K204A, L207A, H208A mutant (Mut) or EMC9 WT. Total cell lysates were analyzed by western blotting with the indicated antibodies. Note that all variants rescue EMC assembly as judged by the levels of EMC8’s partner subunit EMC2. **G)** As in F but in stable RPE1 Ca_v_1.2α•Ca_v_β_3_ reporter cells, Ca_v_1.2α•Ca_v_β_3_•Ca_v_α_2_δ_1_ reporter cells or RPE1 cells transiently transduced with the indicated fluorescent reporters. The GFP/RFP or RFP/GFP ratios of BFP^+^ cells were determined by flow cytometry and are depicted as histograms. **H)** 72h scrambled or EMC1 siRNA knockdown in stable RPE1 Ca_v_1.2α•Ca_v_β_3_ reporter cells. 24h after siRNA transfection, cells were transduced with lenti-viral rescue constructs encoding either just BFP or BFP separated by a 2A site from the prolactin signal sequence (Prl(ss))-ALFA-tagged EMC1 WT (23-end) or EMC1 D961A, R981L mutant (Mut) (23-end). Total cell lysates were analyzed by western blotting with the indicated antibodies. Note that EMC1(Mut) rescues EMC assembly as judged by the levels of EMC core subunit EMC3. **I)** As in H but in RPE1 cells transiently transduced with the indicated fluorescent reporters. The RFP/GFP ratios of the BFP^+^ cells were determined by flow cytometry and are depicted as histograms.

**Figure S3.**
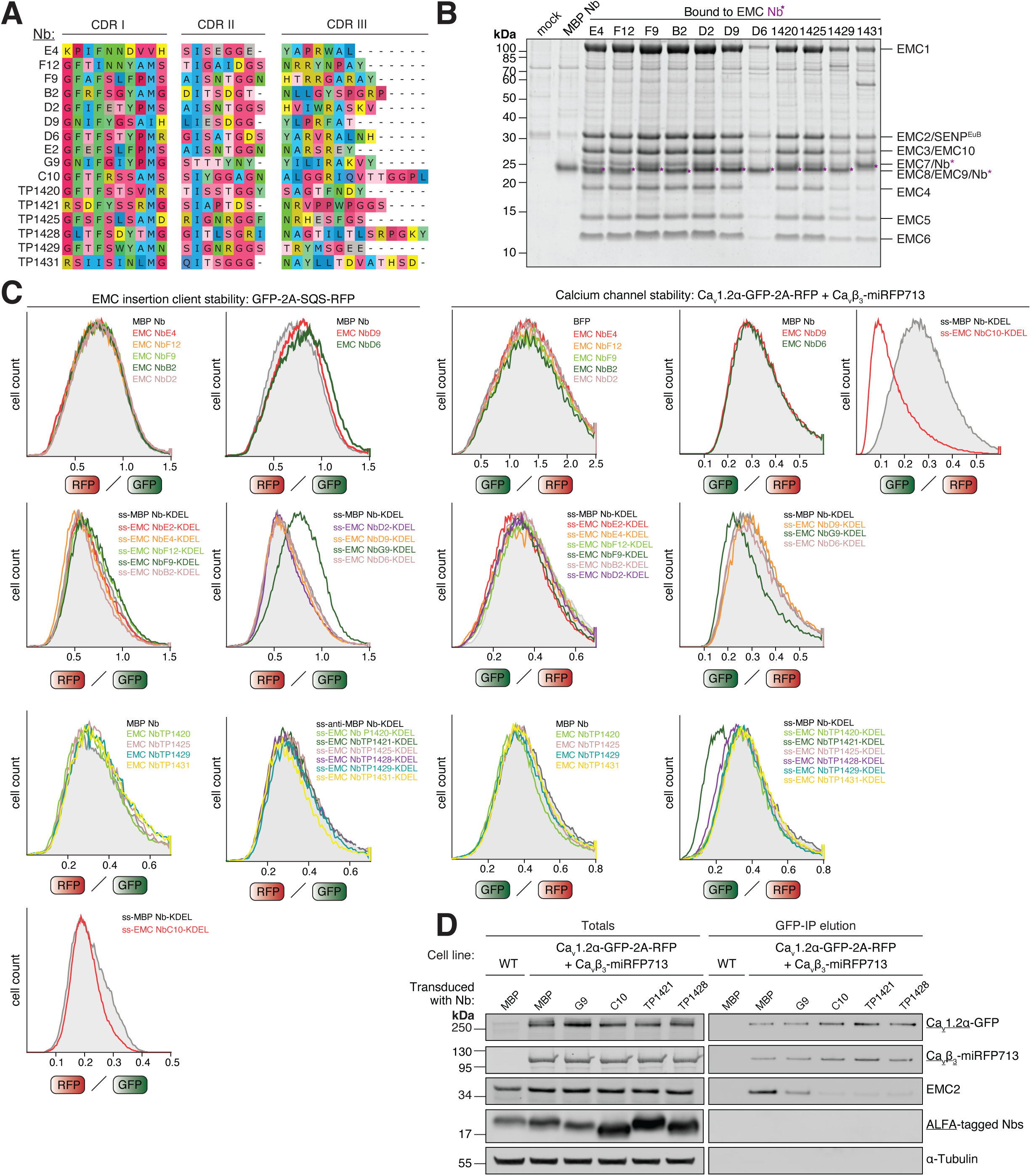
Characterization of the anti-EMC nanobody toolbox. **A)** Focused sequence alignment of the 16 different anti-EMC nanobody (Nb) classes characterized in this study. Only the sequence stretches encompassing the complementarity-determining regions (CDRs) I-III, which constitute the antigen-binding loops of these Nbs, are depicted. **B)** ALFA-GFP-SUMO^Eu^-tagged anti-MBP control or anti-EMC Nbs were immobilized via a biotinylated, SUMOstar protease-cleavable anti-ALFA tag Nb on Streptavidin magnetic beads. The beads were incubated with a GDN-solubilized Expi293 total cell lysate. After washing, proteins bound to the control or anti-EMC Nbs were specifically eluted by native SENP^EuB^ cleavage. The eluate was analyzed by SDS-PAGE and SYPRO Ruby staining. Note that all anti-EMC Nbs specifically purified the nine EMC subunits. Nbs D6, TP1429 and TP1431 had lower affinity and isolated less EMC. **C)** Flow cytometry assay in stable RPE1 Ca_v_1.2 α•Ca_v_β_3_ reporter cells or RPE1 cells transiently transduced with EMC-dependent insertase client reporter SQS. These cell lines were transduced to express the indicated Nbs either from a BFP-2A-Nb cassette in the cytosol or from a BFP-2A-Prl(ss)-Nb-KDEL cassette in the ER lumen. The GFP/RFP or RFP/GFP ratios of BFP^+^ cells are depicted as histograms. Note, that inhibitory Nbs G9, C10, TP1421 and TP1428 also inhibited chaperone function when targeted to the ER lumen. We believe this occurs because small, fast-folding proteins like Nbs might occasionally escape translocation into the ER lumen and accumulate at sufficient levels in the cytosol to cause the observed inhibitory effect. **D)** Stable RPE1 Ca_v_1.2α•Ca_v_β_3_ reporter cells were transduced to express the indicated anti-MBP control or inhibitory anti-EMC Nbs in the cytosol. Cells were lysed in detergent and subjected to anti-GFP Nb purification of Ca_v_1.2α-GFP to assess EMC co-purification in the presence of these Nbs. Total cell lysates and protease elution of the GFP-IP were analyzed by western blotting with the indicated antibodies.

**Figure S4.**
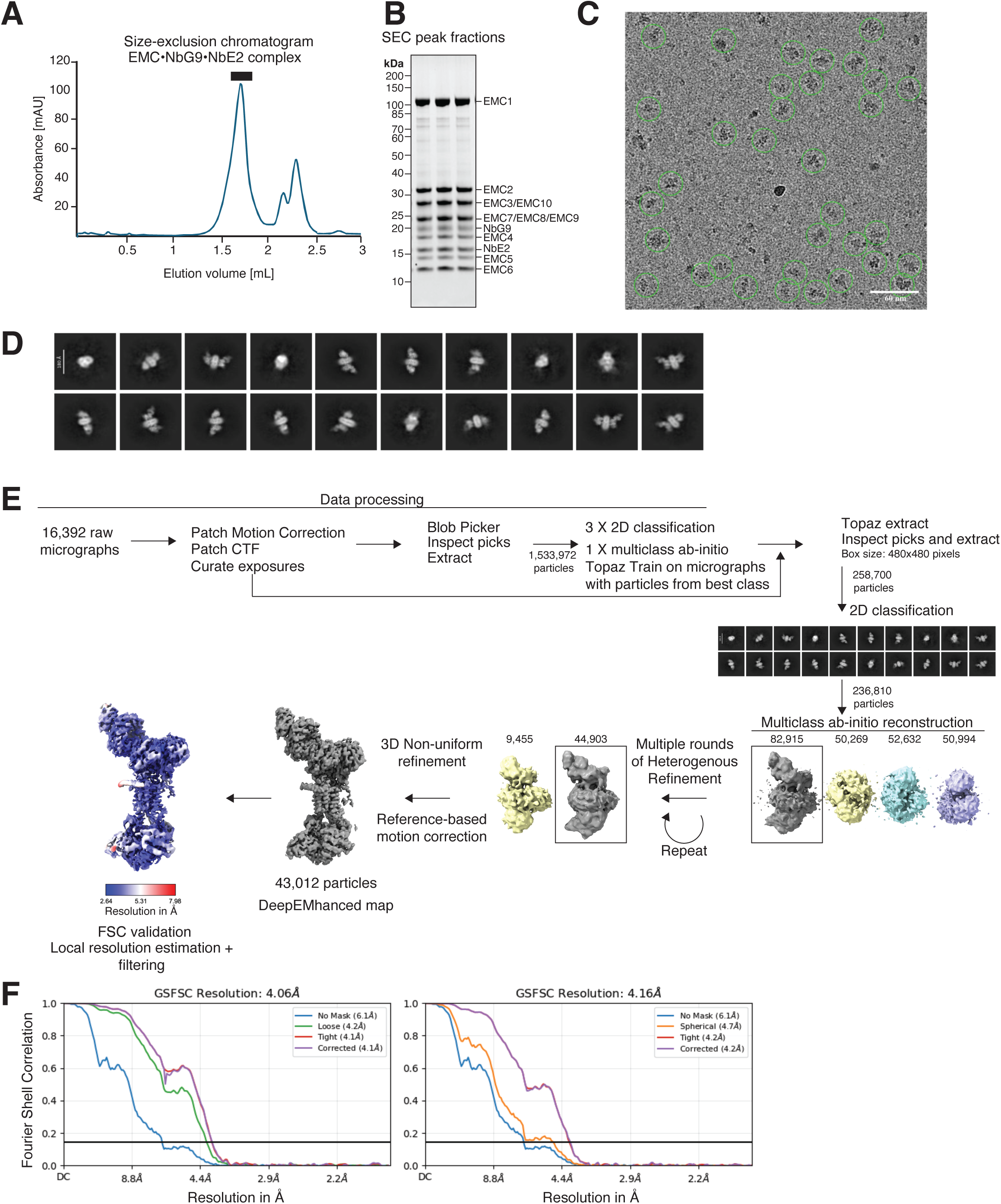
Cryo-EM structure of the human EMC bound to two nanobodies. **A)** Endogenous human EMC was purified from Expi293 cell lysate using biotinylated, SENP^EuB^-cleavable anti-EMC NbE2 and then mixed with purified inhibitory anti-EMC NbG9. The resulting complex was further purified by size-exclusion chromatography (SEC) on a Superose 6 Increase 3.2/300 column. **B)** Analysis of the SEC peak fractions from A by SDS-PAGE and SYPRO Ruby staining. **C)** Representative cryoEM micrograph collected using Titan Krios G2 operating at 300KeV on Falcon 4i detector and SelctrisX energy filter. Particles representing the EMC•NbG9•NbE2 complex picked using manual picker in cryoSPARC are highlighted. **D)** Representative 2D class averages generated during data processing. **E)** General cryoEM data processing workflow employed for obtaining the structure of the EMC in complex with Nbs G9 and E2. The local resolution map was calculated using cryoSPARC local resolution estimation and local resolution filtering methods and ranges from 2.6 Å to 7.9 Å resolution. **F)** Fourier Shell Correlation (FSC) of the final 4.06 Å cryo-EM map of the EMC•NbG9•NbE2 complex with different masks from the Non-uniform refinement and FSC validation, respectively.

**Figure S5.**
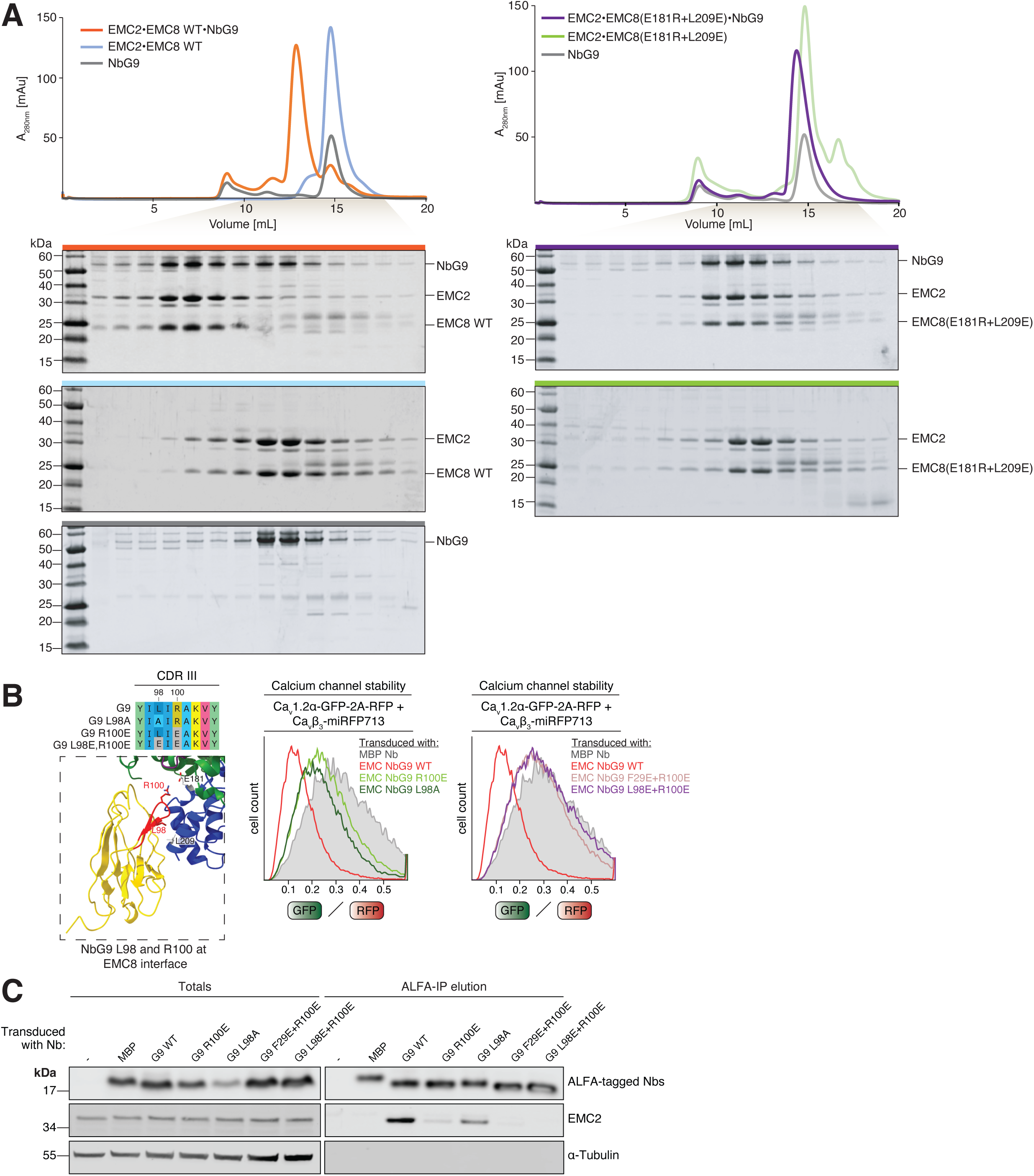
Validation of the EMC•NbG9 cryo-EM structure. **A)** (*Left*) Superdex 200 10/300 size exclusion chromatograms of purified GFP-SUMO^Eu^-tagged NbG9 (grey), purified EMC2•EMC8 complex (blue) or EMC2•EMC8•NbG9 complex (orange) are shown. Peak fractions were analyzed by SDS-PAGE and SYPRO Ruby staining. Gels are color-coded to match their respective chromatography runs. Note that NbG9 binds to the purified EMC2•EMC8 complex and results in a left shift to earlier elution volume, consistent with complex formation. (*Right*) As in A, but with an EMC8 variant that contains two mutations at the EMC8•Nb G9 interface. These mutations abolish ternary complex formation, validating the modeled NbG9 interface on the EMC in our cryo-EM structure. **B)** (*Left*) View of the NbG9•EMC8 interface in our cryo-EM model, highlighting CDR III in red and depicting residues L98 and R100 as sticks. The sequence of NbG9’s CDR III and the mutations made below are highlighted on top. EMC8 residues E181 and L209 mutated in A) are highlighted as light grey sticks. (*Right*) Stable RPE1 Ca_v_1.2α•Ca_v_β_3_ reporter cells were transduced to express either anti-MBP control Nb, wild-type NbG9 or the indicated single and double mutants of NbG9 from a BFP-2A-Nb cassette in the cytosol. The GFP/RFP ratios of BFP^+^ cells are depicted as histograms. F29 is located in CDR I. Note that mutations of NbG9 CDR residues abolish its inhibitory effect. **C)** RPE1 cells transduced with the indicated, ALFA-tagged Nbs were lysed in detergent and subjected to anti-ALFA tag Nb purification. Total cell lysates and ALFA Nb protease eluates were analyzed by western blotting with the indicated antibodies. Note that disruption of NbG9’s CDR residues reduces its ability to interact with the EMC.

**Figure S6.**
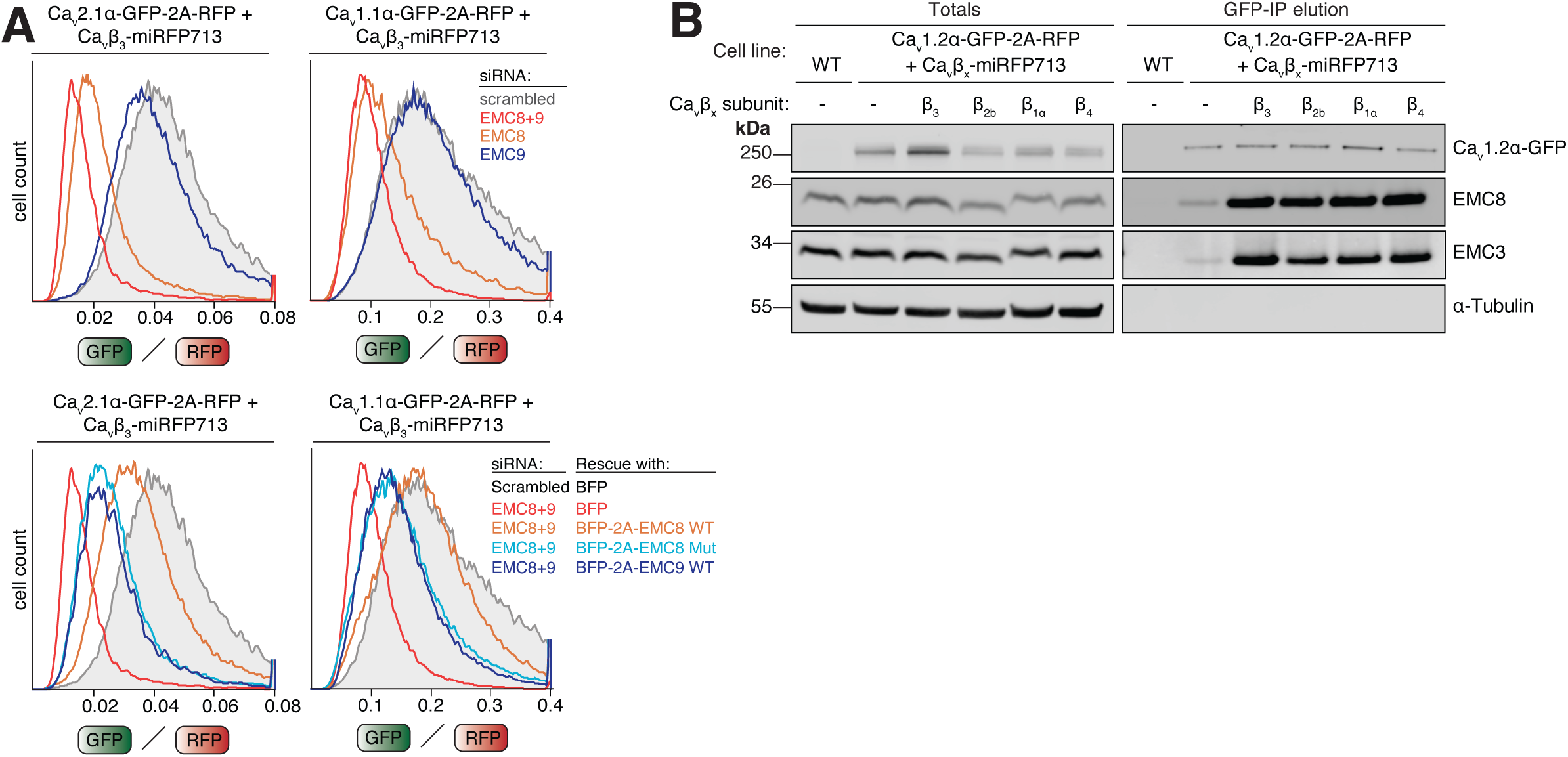
Biogenesis of brain and skeletal muscle calcium channels also relies on EMC chaperone function. **A)** Experiments in stable RPE1 cells expressing the brain Ca_v_2.1α•Ca_v_β_3_ channel (*left*) or skeletal muscle Ca_v_1.1α •Ca_v_β_3_ channel reporter (*right*). (*Top*) 72h scrambled, EMC8, EMC9 or EMC8+9 double siRNA knockdown. The GFP/RFP ratios of the resulting cells were determined by flow cytometry and are depicted as histograms. (*Bottom*) 72h scrambled or EMC8+9 double siRNA knockdown. 24h after siRNA transfection, cells were transduced with lenti-viral rescue constructs encoding either just BFP or BFP separated by a 2A site from either EMC8 WT, EMC8 K204A, L207A, H208A mutant (Mut) or EMC9 WT. The GFP/RFP ratios of BFP^+^ cells were determined by flow cytometry and are depicted as histograms. **B)** Stable RPE1 Ca_v_1.2α reporter cells expressing the indicated Ca_v_β subunits fused to miRFP713 were lysed in detergent and subjected to anti-GFP Nb purification of Ca_v_1.2α-GFP to assess EMC co-purification in the presence of the different Ca_v_β subunits. Total cell lysates and protease elution of the GFP-IP were analyzed by western blotting with the indicated antibodies.

**Figure S7.**
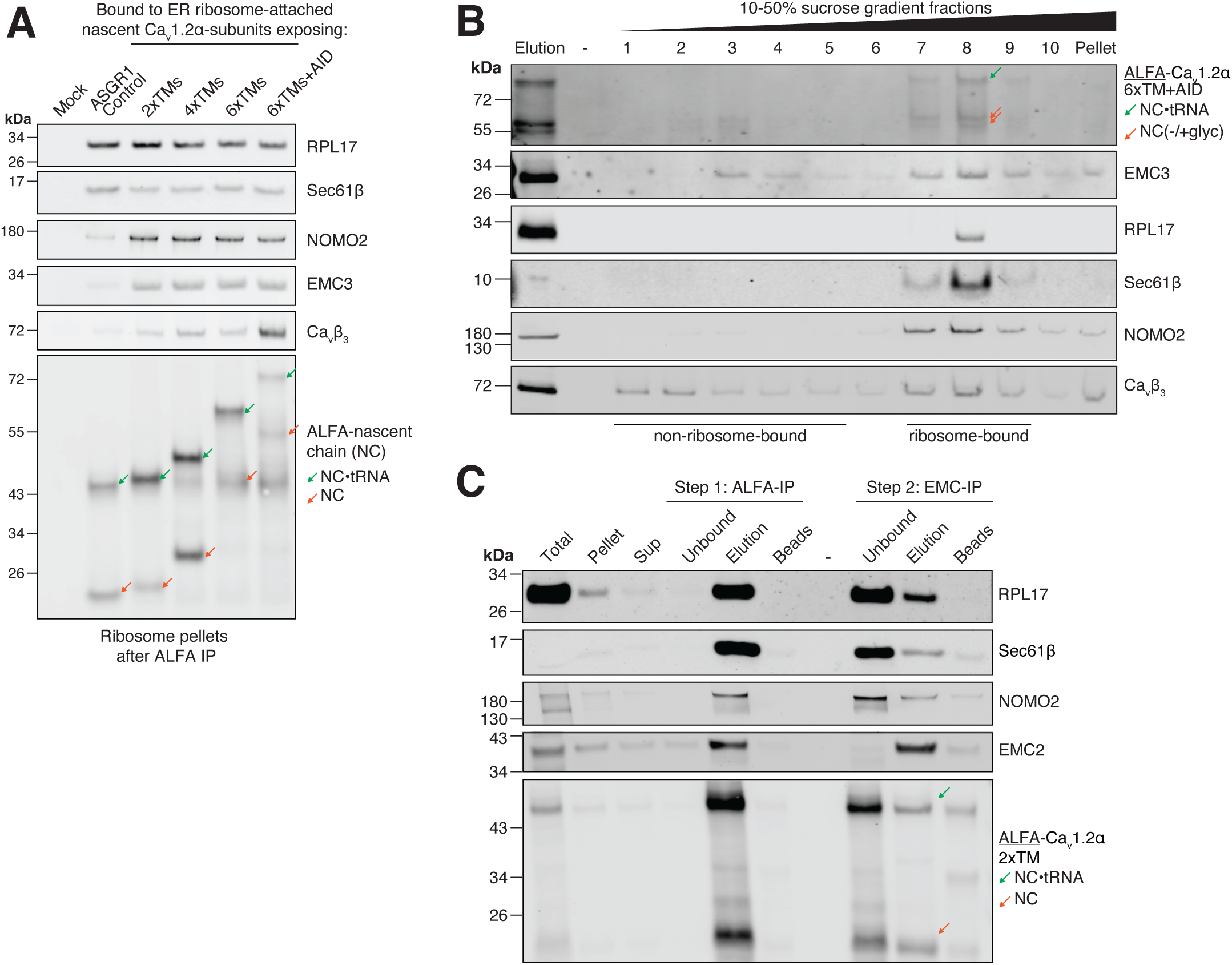
The EMC associates co-translationally with Ca_v_1.2α-synthesizing ribosome•MPT complexes. **A)** Assay as in Fig. 5C. Normalized ribosome pellets of ribosome-nascent chain complexes (RNCs) stalled while translating either an EMC-independent single-pass control protein ASGR1, or Ca_v_1.2α exposing 2x, 4x, 6x or 6xTMs plus the AID-containing cytosolic linker between bundles I and II outside of the ribosome exit tunnel, were analyzed by western blotting with the indicated antibodies. A mock translation without RNA served as a background control. Note that EMC engagement with TM bundle I intermediates of Ca_v_1.2α is constant. **B)** Stalled RNCs containing nascent Ca_v_1.2α exposing 6xTMs+AID were purified by ALFA nanobody IP as described in Fig. 5B and the eluate was then overlayed onto a 10-50% sucrose gradient. After the run, eleven fractions were collected from the top of the gradient for analysis by western blotting. Note that ribosomes and ribosome-bound proteins peak around fraction 8. Both EMC and Ca_v_β_3_ were found to migrate with ribosomes in the sucrose gradient. **C)** Stalled RNCs containing Ca_v_1.2α exposing 2xTMs were generated as described in Fig. 5B. After purification of ALFA-tagged RNCs using a SUMOstar-cleavable anti-ALFA-nanobody, the resulting protease eluate was subjected to a second IP with inert, SENP^EuB^-cleavable anti-EMC NbE2. Samples of this 2-step IP were analyzed by western blotting with the indicated antibodies. Note that the EMC is bound to intact RNCs containing tRNA-associated nascent chain (NC), ribosomes (RPL17), the translocon (Sec61β) and MPT BOS complex subunit NOMO2.

**Table S1.**
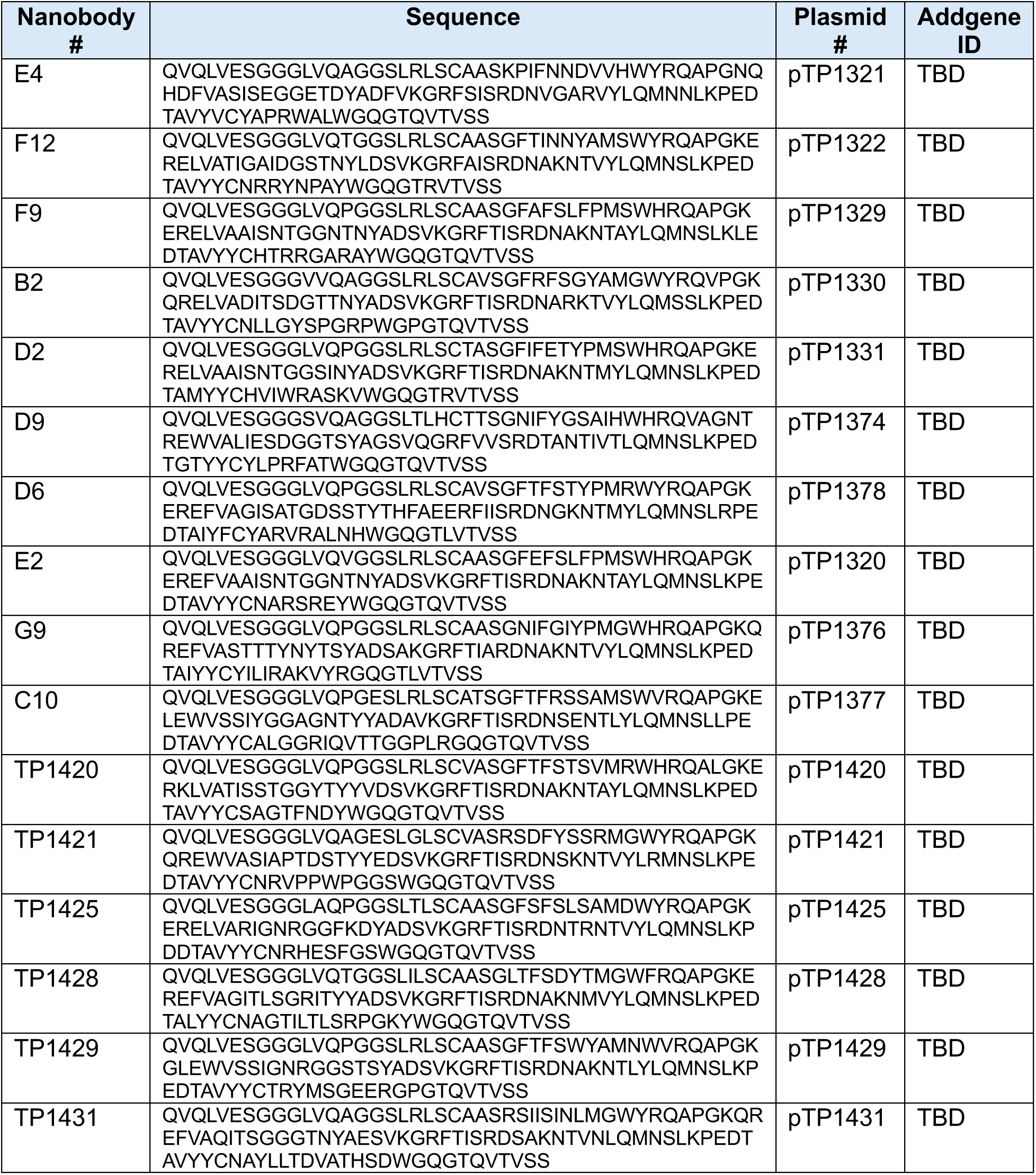
Protein sequences of anti-EMC nanobodies generated in this study.

**Table S2.**
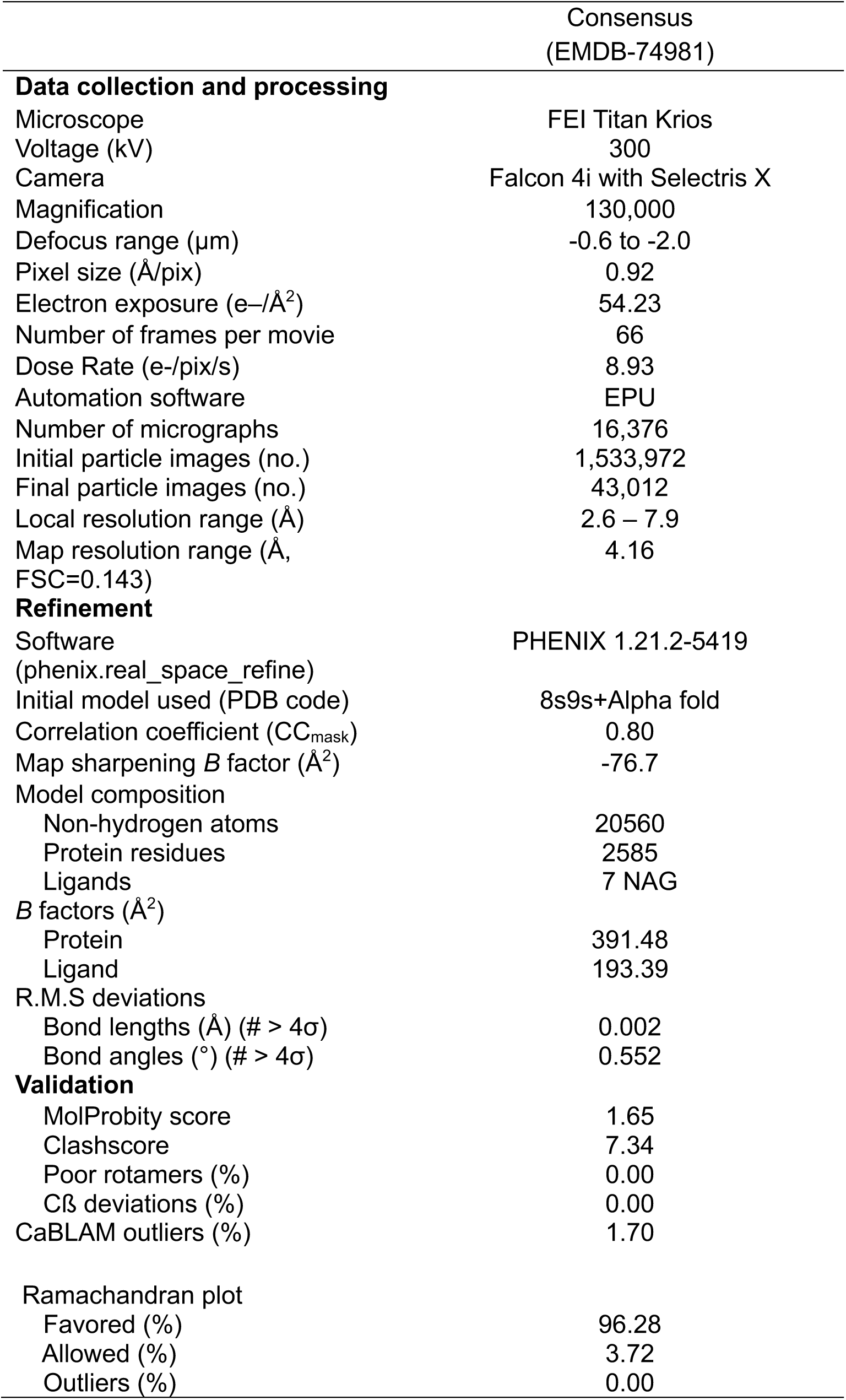
Cryo-EM data collection, refinement, and validation statistics.

## Additional supplemental items

**Supplemental Video S1. Live cell imaging of calcium dynamics in cardiomyocytes transduced with control anti-MBP nanobody.** Cytosolic BFP is expressed from the same lenti-viral vector as the nanobody and served as a transduction marker. GCaMP6f fluorescence in the green channel indicates calcium dynamics accompanying cardiomyocyte contraction.

**Supplemental Video S2. Live cell imaging of calcium dynamics in cardiomyocytes transduced with control anti-EMC nanobody G9.** Cytosolic BFP is expressed from the same lenti-viral vector as the nanobody and served as a transduction marker. GCaMP6f fluorescence in the green channel indicates calcium dynamics accompanying cardiomyocyte contraction.

**Supplemental Video S3. Live cell imaging of calcium dynamics in cardiomyocytes transduced with control anti-EMC nanobody C10.** Cytosolic BFP is expressed from the same lenti-viral vector as the nanobody and served as a transduction marker. GCaMP6f fluorescence in the green channel indicates calcium dynamics accompanying cardiomyocyte contraction.

